# Molecular insights into the interaction between a disordered protein and a folded RNA

**DOI:** 10.1101/2024.06.12.598678

**Authors:** Rishav Mitra, Emery T. Usher, Selin Dedeoğlu, Matthew J. Crotteau, Olivia A. Fraser, Neela H. Yennawar, Varun V. Gadkari, Brandon T. Ruotolo, Alex S. Holehouse, Loïc Salmon, Scott A. Showalter, James C. A. Bardwell

**Affiliations:** Howard Hughes Medical Institute, University of Michigan, Ann Arbor, MI 48109, USA; Department of Molecular, Cellular, and Developmental Biology, University of Michigan, Ann Arbor, MI 48109, USA; Department of Biochemistry and Molecular Biophysics, Washington University School of Medicine, St. Louis, MO, USA; Center for Biomolecular Condensates (CBC), Washington University in St. Louis, St. Louis, MO, USA; Centre de Résonance Magnétique Nucléaire à Très Hauts Champs, (CRMN), UMR 5082, CNRS, ENS Lyon, UCBL, Université de Lyon, 69100 Villeurbanne, France; Center for Eukaryotic Gene Regulation, Department of Biochemistry and Molecular Biology, The Pennsylvania State University, University Park, PA 16802, USA; The Huck Institutes of the Life Sciences, The Pennsylvania State University, University Park, PA 16802, USA; Department of Chemistry, The Pennsylvania State University, University Park, PA 16802, USA; Department of Chemistry, University of Michigan, Ann Arbor, MI 48109, USA

**Author notes:** Department of Bioengineering and Therapeutic Sciences, University of California San Francisco, San Francisco, CA 94158, USA. Department of Chemistry, University of Minnesota, Minneapolis, MN 55455, USA. These authors contributed equally (Rishav Mitra, Emery T. Usher).

**Keywords:** intrinsically disordered regions (IDRs), phase separatio, | condensate, RNA-binding protein, RNA-protein complex, nuclear magnetic resonance (NMR)

## Abstract

Intrinsically disordered protein regions (IDRs) are well-established as contributors to intermolecular interactions and the formation of biomolecular condensates. In particular, RNA-binding proteins (RBPs) often harbor IDRs in addition to folded RNA-binding domains that contribute to RBP function. To understand the dynamic interactions of an IDR-RNA complex, we characterized the RNA-binding features of a small (68 residues), positively charged IDR-containing protein, SERF. At high concentrations, SERF and RNA undergo charge-driven associative phase separation to form a protein- and RNA-rich dense phase. A key advantage of this model system is that this threshold for demixing is sufficiently high that we could use solution-state biophysical methods to interrogate the stoichiometric complexes of SERF with RNA in the one-phase regime. Herein, we describe our comprehensive characterization of SERF alone and in complex with a small fragment of the HIV-1 TAR RNA (TAR) with complementary biophysical methods and molecular simulations. We find that this binding event is not accompanied by the acquisition of structure by either molecule; however, we see evidence for a modest global compaction of the SERF ensemble when bound to RNA. This behavior likely reflects attenuated charge repulsion within SERF via binding to the polyanionic RNA and provides a rationale for the higher-order assembly of SERF in the context of RNA. We envision that the SERF-RNA system will lower the barrier to accessing the details that support IDR-RNA interactions and likewise deepen our understanding of the role of IDR-RNA contacts in complex formation and liquid-liquid phase separation.

**SIGNIFICANCE:** Subcellular organization through the formation of biomolecular condensates has emerged as an important contributor to myriad cellular functions, with implications in homeostasis, stress response, and disease. To understand the general and specific principles that support condensate formation, we must interrogate the interactions and assembly of their constituent biomolecules. To this end, this study introduces a simple model system comprised of a small, disordered protein and small RNA that undergo charge-driven, associative phase separation. In addition to extensive biophysical characterization of these molecules and their complex, we also generate new insights into mode of interaction and assembly between an unstructured protein and a structured RNA.

## INTRODUCTION

Intermolecular interactions underscore many critical functions as a means to interpret, transmit, and respond to numerous and diverse signals in a cell. Among the proteins involved in this so-called ‘information processing’, many harbor intrinsically disordered regions (IDRs) that mediate interactions even though they lack stable tertiary structures^1^. Although they have roles in virtually every cellular process, IDRs have emerged as important determinants for proteins binding to RNA^2–4^. Previously, RNA-binding proteins (RBPs) were assigned as such based on the presence of a structured RNA-binding domain ^5^(5), but recent large-scale interactome studies illuminated many ‘unconventional’ RBPs with IDRs that can directly engage RNA^6–8^. Such RBPs may leverage both folded and disordered RNA-interaction modules in concert to bind RNA with tailored affinity or specificity^3^. Moreover, IDR-mediated interactions are not limited to binary complexes; many IDR-containing RBPs undergo higher order assembly with RNA to form biomolecular condensates^9^.

Biomolecular condensates are non-stoichiometric assemblies of biomolecules that concentrate specific components while excluding others^9^. These multicomponent cellular bodies play key roles in the spatiotemporal regulation of biological processes in space and time^9,10^.

While condensates can form through various mechanisms, the physics of phase separation offers a parsimonious explanation for their formation, behavior, and dissolution^11–13^. Condensates often contain protein and RNA, and both can play key roles in determining condensate formation and function^14–17^. The importance of IDRs and RNA in the context of condensate stems from their inherent tendency to form multivalent interactions ^18^. This is especially true for polycationic disordered regions, which interact directly with polyanionic RNA^19–22^. As such, charge-mediated RNA-protein interactions are central in determining condensate assembly, regulation, and disassembly. So far, high-resolution biophysical investigation into the intermolecular interactions between IDRs and RNA molecules that underlie condensate formation have largely been limited to artificial peptides and simple homopolymeric RNA species which have provided simple, though not particularly physiological systems to elucidate basic physical principles^20–25^.

Several practical and technical barriers have historically impeded high-resolution, quantitative characterization of IDR-RNA complexes. First, the structural heterogeneity of IDRs (and some RNAs) generally precludes classical structural interrogation methods (i.e., X-ray crystallography); therefore, we must rely on solution-state ensemble methods. Second, many of the ‘classical’ RBPs, such as FUS, hnRNPA1, and G3BP1, are large, often aggregation-prone, and have multi-domain architecture, such that the residue-level details of RNA binding are challenging to obtain^26–28^. Similarly, endogenous RNA molecules can be kilobases in length, which can pose a challenge to accessing high-resolution structural and dynamic information of the molecule^9^. Finally, at the intersection of the two aforementioned barriers, many of the techniques used to characterize IDRs and their complexes, such as nuclear magnetic resonance (NMR) spectroscopy or small angle X-ray scattering (SAXS), require relatively high concentrations of biomolecules. For large, multi-domain RBPs, such a concentration regime may be inaccessible for RBP-RNA partners that readily de-mix and, hence, preclude the study of stoichiometric complexes.

In this work, we sought to circumvent these challenges by investigating a pair of biomolecules in which IDR and RNA properties in the bound and unbound states could be directly interrogated by experiments and simulations. The ideal IDR-RNA system should undergo phase separation at sufficiently high concentrations such that the stoichiometric complex can be studied in the absence of condensate formation. In addition, both molecules should be relatively small (i.e., single domain) in order to make spectroscopic interpretation more straightforward. To this end, we selected the 68-residue, intrinsically disordered *Saccharomyces cerevisiae* protein, SERF (UniProt ID: YDL085C-A) (**Fig. 1A**). SERF (<u>S</u>mall <u>E</u>RDK-<u>R</u>ich <u>F</u>actor) is highly charged with a bias toward positive residues and is predicted to be primarily intrinsically disordered (**Fig. 1B**). Structural studies of the *Caenorhabditis elegans* SERF homolog, MOAG-4, showed that the C-terminal region of the protein adopts an α-helix, whereas the N-terminal region is more structurally heterogeneous^29^. Recent work showed that the human SERF protein partitions to nucleoli and can facilitate the nucleolar incorporation of fluorescently labeled RNA^30^. For the minimal RNA component, we selected the apical region of the Trans-Activation Response element (TAR) RNA from human immunodeficiency virus type-1 (HIV-1). TAR is among the best-studied helix-junction-helix motifs and, importantly, moves us beyond the study of homopolymeric RNAs^31,32^. This 29-nucleotide RNA fragment (depicted in **Fig. 1C**) contains two A-form helical stems (Helix I and Helix II), a trinucleotide bulge, and a hexanucleotide apical loop, which collectively undergo complex dynamic motions over a broad range of timescales^32^. Extensive prior research into TAR and its interactions in the context of virulence equips us with a wealth of information about its structural and dynamic properties^33–35^.

**Figure 1:**
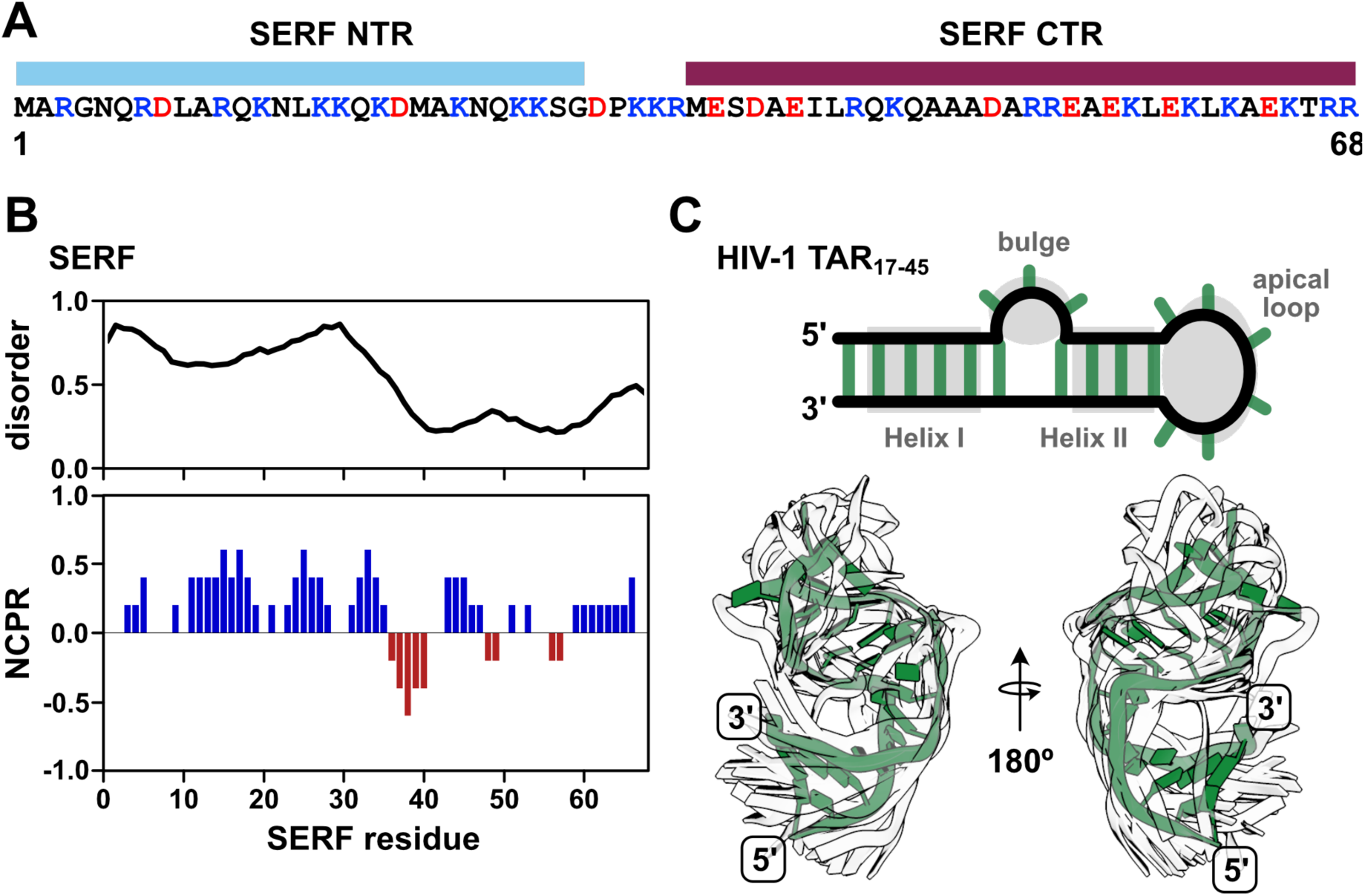
SERF is a highly positively charged disordered protein that can bind RNA. **(A)** Sequence of *S. cerevisiae* SERF with domain annotations (NTR – N-terminal region; CTR – Cterminal region). Positive and negative amino acids are colored in blue and red, respectively. **(B)** Per-residue disorder scores from Metapredict(1) (top) and SERF net charge per residue (NCPR) calculated with CIDER using a window size of five(2). **(C)** Diagram of the fragment of the HIV-1 TAR RNA used in this study (nucleotides 17-45) (top). TAR features of interest are shown in gray. Helix I and Helix II are also referenced as ‘upper’ and ‘lower’ helices, respectively. The bottom panel shows the alignment of TAR conformers from the NMR solution structure (PDB ID: 1ANR)(3). One conformer is shown in green for clarity; the other 19 are shown as white outlines to illustrate TAR dynamics.

Our work establishes that SERF and TAR primarily form a binary complex at concentrations suitable for biophysical characterization (< 0.5 mM) and can form biomolecular condensates *in vitro* beyond this point or by adding crowding agents. In the SERF:TAR complex, neither SERF nor TAR undergo major structural rearrangements; SERF retains its conformational flexibility and TAR similarly retains its base-paired secondary structure.

Through extensive biophysical characterization of the protein, the RNA, and their complex, this study sheds light on the driving forces behind the interactions of a disordered protein with a folded RNA that may be extended to more complicated, multi-domain RBPs.

## RESULTS

### Conformational properties of SERF

Prior to investigating the molecular mechanisms of SERF-RNA interaction, we first characterized the SERF protein in isolation using multiple biophysical methods. We used solution NMR spectroscopy to obtain residue-level information on the structural and dynamic features of the SERF conformational ensemble. Like many disordered proteins, SERF shows limited signal dispersion in its ^1^H,^15^N-HSQC spectrum (**Fig. S1A**), which poses a challenge for resonance assignments and subsequent structural characterization^36^. Therefore, we turned to ^13^C direct-detected NMR spectroscopy, which offers improved resolution and reduced peak overlap for IDRs compared to proton-based detection^37,38^. The ^13^C, ^15^N-CON spectrum of SERF shows excellent chemical shift dispersion and enabled us to obtain unambiguous backbone and sidechain assignments for each SERF residue (**Fig. S1B, Table S1**).

From the disorder prediction profile for *S. cerevisiae* SERF (**Fig. 1B**) and from structural studies on other SERF family proteins, we expected to observe some structural features in the C-terminal region of SERF (CTR). To quantify possible secondary structural features of SERF, we calculated per-residue secondary structure propensities from the deviations of the measured C_α_ and C_β_ chemical shifts referenced against theoretical SERF chemical shifts assuming a random coil^39^. We found that the CTR exhibits α-helical character between residues 36 and 60 (**Fig. 2A**). The N-terminal region of SERF (NTR) lacks secondary structure aside from a short α-helical stretch near residues 5-15; the differences in magnitude between the N- and C-terminal helical regions suggest that the helix in the NTR is less populated relative to the helix in the CTR (**Fig. 2A**).

**Figure 2:**
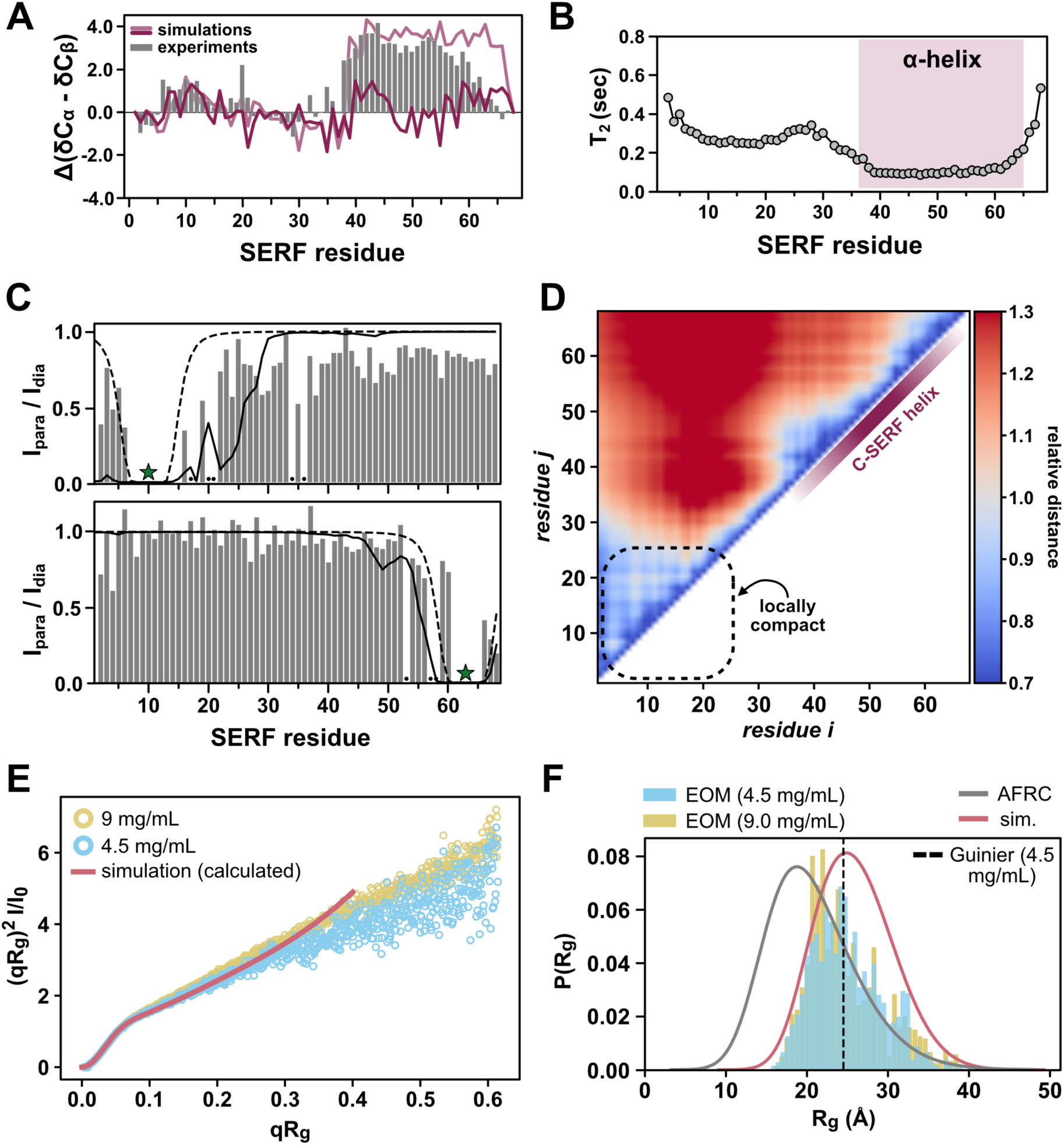
SERF is globally disordered amid regions of transient structure. **(A)** Secondary structure of SERF calculated from NMR chemical shifts (grey bars). Positive values indicate regions of alpha helix; negative values correspond to beta strand/extended regions. Average alpha helix character of SERF determined using back-calculated chemical shifts from all-atom simulations is shown as a dark pink trace. For comparison, back-calculated chemical shifts from simulations in which C-terminal region of SERF was fixed as a helix are shown in light pink. **(B)** Residue-specific ^15^N transverse relaxation times (*T_2_*) of SERF. The C-terminal helix (shaded) corresponds to a track of residues with depressed T_2_ values. **(C)** Paramagnetic relaxation enhancement (PRE) NMR profiles for SERF containing PRE labels at residues 10 (top) or 63 (bottom). Grey bars reflect PRE measurements: missing/unassignable resonances are denoted with a black dot. The solid line is the PRE profile calculated from all-atom simulations and the dashed line is the theoretical profile for a Flory Random Coil (null model). The star indicates the site of MTSSL attachment in PRE NMR experiments.

To further explore the local and global features of the SERF ensemble, we performed all-atom Monte Carlo simulations of SERF using the CAMPARI/ABSINTH simulation package with OPLS-AA/L force field parameters^40^. This simulation strategy has been applied to several disordered protein systems with good agreement with *in vitro* measurements^41–44^. We observe qualitative agreement between the experimental secondary structure profile and that back calculated from SERF simulations (see *Methods*) (**Fig. 2A, Fig. S1C-D**). The simulations capture the modest helical region in NTR but fail to reproduce the more populated helix in the CTR. SERF simulations show high helical character near residues 40-45 and toward the distal C-terminus (residues 55-65), but the intervening stretch of residues is much less helical compared to experiments (**Fig. 2A, Fig. S2A**). Possible explanations for this discrepancy are detailed in the simulation *Methods*.

To evaluate the local conformational properties of SERF, we next measured ^15^N spin relaxation using ^13^C direct-detect NMR methods optimized for IDRs^45^. The T_1_ and T_2_ relaxation times are sensitive to both global chain reorientation and local backbone motions that occur on the picosecond-nanosecond timescale^46^. The T_1_ profile for SERF is largely featureless with relaxation times typical for a small IDR (**Fig. S1E**)^45^. Fast molecular rearrangements, like those that are characteristic for unstructured protein regions, generally confer longer T_2_ times as compared to more rigid, structured regions. Compared to the NTR, the T_2_ values in the CTR of SERF are depressed (**Fig. 2B**), likely reflecting the high contact density between spin systems arising from the α-helical structure. The marked differences in T_2_ times between the two halves of SERF suggest that the molecular motions of these regions are decorrelated (**Fig. 2B**).

To complement residue-level insights into the SERF ensemble, we next interrogated the global conformational properties of SERF. Despite the lack of a stable tertiary structure, transient tertiary contacts and sparsely sampled compact conformations often contribute to IDR ensembles ^47,48^. Thus, we asked whether transient tertiary contacts between the N- and C-terminal regions of SERF contribute to its ensemble behavior. Paramagnetic relaxation enhancement (PRE) NMR methods are highly sensitive to sparsely populated conformations with long-range contacts and can provide site-resolved information about such interactions^49^. We labeled one of two exogenous cysteines on SERF at either residue 10 or 63 with the nitroxide spin label, (1-Oxyl-2,2,5,5-tetramethylpyrroline-3-methyl) methanethiosulfonate (MTSSL), and then measured PRE using ^13^C direct-detect NMR experiments. In this method, resonances for residues within ∼20 Å labeled residue are expected to decrease in intensity due to enhanced relaxation via the unpaired electron on MTSSL^50^.

Our PRE results, presented in **Fig. 2C** as the ratio of peak intensities in the paramagnetic to diamagnetic measurements, suggest minimal long-range interactions in the ensemble; rather, PRE occurs almost exclusively around the site of labeling (**Fig. 2C**). This conclusion is supported by PRE profiles calculated from all-atom simulations of SERF. Finally, distance maps determined from simulations quantify ensemble-averaged inter-residue distances, normalized by the distance expected for a random coil (**Fig. 2D**) (see *Methods*). The distance map reflects the modest relative compaction in the NTR that we also observed in the experimental PRE profile for this region (**Fig. 2C**) and similarly suggests that SERF lacks long-range intermolecular contacts. Collectively, the spin relaxation measurements, PRE experiments, and all-atom simulations show that SERF is primarily unstructured, but contains elements of local structure and/or compactness that bias its global ensemble away from that of a random coil.

We next used small-angle X-ray scattering (SAXS) to quantify the global ensemble properties of SERF. SAXS reports on the size (reported as radius of gyration (R_g_)) and shape of a molecule based on the features of the scattering pattern ^51^. The scattering profiles obtained from two different concentrations of SERF are consistent with a primarily disordered protein. Further, the scattering profile calculated from all-atom simulations shows excellent agreement with both experimental datasets (**Fig. 2E**, **Fig. S2**). We determined the ensemble-averaged R_g_ using the Guinier approximation, ensemble optimization method (EOM), and the Molecular Form Factor (MFF) of Riback et al. (see *Methods*)^52–54^. These different approaches showed good agreement with each other, with average R_g_ values of 24.5 ± 0.2 Å (Guinier), 25.2 Å (EOM), and 23.9 ± 0.1 Å (MFF). These are also in good agreement with R_g_ values calculated from all-atom simulations (R_g_ = 25.9 Å) and from the deep learning-based predictor, ALBATROSS (R_g_ = 24.8 Å) **(Table S2)**^55^. Compared to a null model that assumes random coil polymer behavior, which predicts an average R_g_ of 20.8 Å, the SERF ensemble appears to be somewhat expanded based on both simulations and experiments (**Fig. 2F**)^56^. Deviation from the random coil model likely arises from the high fraction of charged residues in SERF, through which electrostatic repulsion could drive ensemble expansion in a solution with modest ionic strength (such as the buffers used in our experiments). Still, the distribution of sizes that SERF can access is additionally biased by areas of local compactness (i.e., residues 5-25) and secondary structure elements (i.e., helix in the CTR).

### Dynamic SERF-TAR complexes are globally compact

Equipped with a detailed ensemble description of unbound SERF, we next examined the *in vitro* RNA-binding behavior of SERF. An important feature of the SERF-RNA model system is the relatively high concentrations of SERF and RNA required to drive phase separation in the absence of PEG. This enables us to easily work in a concentration regime suitable for biophysical measurements while remaining below the threshold for phase separation. To describe SERF’s dilute-phase interactions with RNA, we measured SERF binding to the bulged stem loop of HIV-1 TAR (nucleotides 19-45, hereafter called ‘TAR’), by fluorescence anisotropy. We monitored the change in fluorescence anisotropy as a function of SERF concentration using fluorescently labeled RNA as the signal source **(Fig. 3A)**. By fitting these data to a 1:1 binding model, we determined that the SERF-TAR complex has an apparent dissociation constant (K_D_) of 0.67 ± 0.04 µM. Equilibrium binding measurements between SERF and fluorescently labeled (rU)_30_ suggest a modestly weaker interaction with this unstructured RNA (K_D_ = 1.9 ± 0.2 µM) as compared to the stem-loop TAR structure (**Table S3**). These values are almost three orders of magnitude weaker than the binding affinity of TAR to its endogenous partner, the Tat protein^57^. This is unsurprising given that Tat-TAR affinity is highly sensitive to mutations to Tat that alter the charge landscape in the Arg-rich binding motif and that the SERF-NTR bears little resemblance to this sequence^58^.

**Figure 3:**
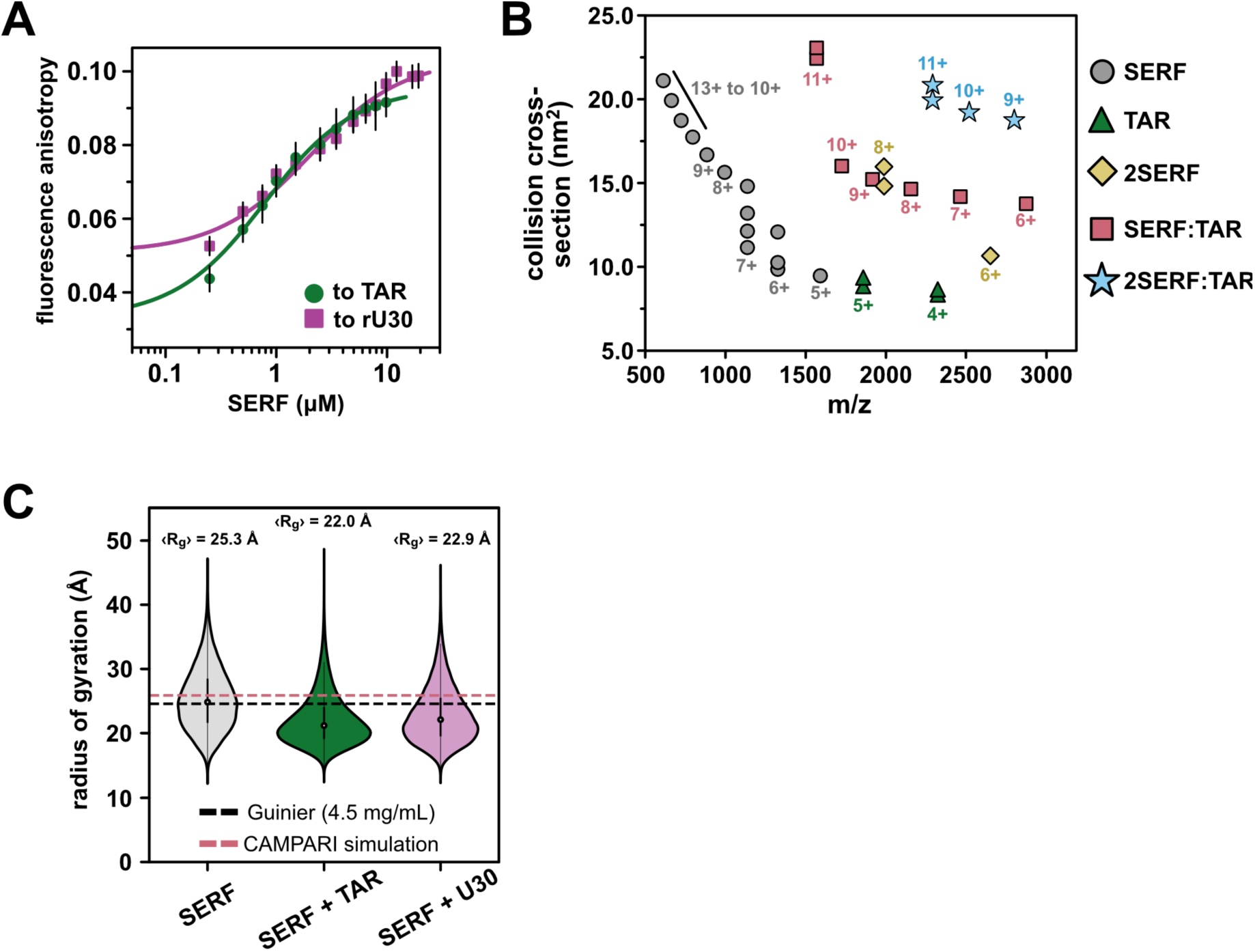
Properties of the SERF-TAR complex. **(A)** Binding isotherms for SERF and FAMlabeled TAR (circles) or FAM-rU30 (squares). The error bars represent the 95% confidence interval from three technical replicates. The solid lines represent the nonlinear fit to a 1:1 binding model. The difference in baseline anisotropy values between binding TAR and rU30 arises from the difference in diffusion properties between the structured vs. unstructured fluorescent RNAs. **(B)** Collision cross section distributions (nm_2_) for the mixture of SERF and TAR determine by IM-MS. **(C)** Distributions of R_g_ values for SERF from coarse-grained simulations of SERF alone (left), SERF and TAR (middle), or SERF and rU30 (right). Note that R_g_ calculations were performed on the protein molecule only. For each case, the average R_g_ is given above its corresponding violin distribution. For comparison, the average R_g_ from all-atom simulations is denoted by the dashed pink line. The R_g_ from Guinier analysis of the SAXS data is shown as the dashed black line. **(D)** Normalized distance map of the ySERF ensemble from simulations. Inter-residue (C_α_-C_α_) distances were measured per frame and averaged; the *relative distance* is presented. Intramolecular distances relative to the AFRC model are presented such that red is more expanded than AFRC and blue is more compact. **(E)** Dimensionless Kratky transformation of scattering data. FoXS was used to calculate the average scattering profile from all-atom simulations.(4) **(F)** Radius of gyration (R_g_) probability distributions for ySERF from SAXS experiments and simulations. The dashed vertical line marks the R_g_ determined from the Guinier approximation. EOM distributions for two different ySERF concentrations are shown as bars colored in blue or yellow. The grey distribution is that of the AFRC model for the ySERF sequence and the pink trace is the distribution of R_g_ values from all-atom ySERF simulations. For comparison, the R_g_ distribution from 14250 frames was normalized by kernel density estimate.

To investigate potential conformational changes coupled to complex formation, we turned to native ion mobility-mass spectrometry (IM-MS). Native IM-MS separates biomolecules based on their shape, surface area, charge, and mass and can be used to determine global molecular dimensions (see *Methods*). Native IM-MS and collision cross section (CCS) measurements have been correlated previously with solution phase fluorescence measurements as well as the computed collision cross section of published crystal structures suggesting that IM-MS can resolve changes in global conformations as small as 3%^59–62^. For ions of the same mass, unfolded or structurally extended species typically have greater CCSs than folded or compact molecules of the same mass. The mass spectrum of an equimolar SERF and TAR mixture shows species corresponding to free SERF (monomer and dimer), free TAR, 1:1 SERF-TAR complex, and a 2:1 SERF-TAR complex **(Fig. 3B, S3A-B)**.

Monomeric SERF yielded a broad charge-state distribution with charge states sampling 5+ to 13+, indicative of an extended conformational ensemble. As determined previously, the CCS distribution for SERF ranges from 9 to 22 nm^2^ (**Table S4**)^63^. SERF dimers were also detected in the IM-MS spectrum^63^. It is possible that ionization in this method reduces intrachain electrostatic repulsion to a sufficient extent to permit SERF-SERF interactions, considering we see no evidence from other methods for SERF dimers. TAR ionizes in primarily two charge states (4+ and 5+) each with a bimodal CCS distribution. Together, across charge states, the IM-MS measurements of TAR reveal a narrow distribution of CCS populations, consistent with the conformational ensemble derived from NMR residual dipolar coupling data^64^. The relatively narrow CCS distribution of the SERF:TAR complexes (from 14 to 22 nm^2^) compared to SERF monomers suggests a compaction of SERF associated with TAR binding.

To probe the structural determinants of SERF-RNA interactions in greater depth, we performed coarse-grained (CG) molecular dynamics simulations to model the SERF-RNA complex using the Mpipi force field^65^. In this model, each amino acid or ribonucleotide is represented by a bead with residue type-specific size and interaction potential^66^. In these simulations, SERF and (rU)_29_ RNA were modeled as flexible polymers, whereas TAR was modeled as a collection of structured conformers using a published TAR NMR ensemble model (PDB ID: 1ANR) (see *Methods* for details). We simulated ‘SERF’ alone or in a box with a structured ‘TAR’ or unstructured ‘(rU)_29_’. For both RNA binding partners, SERF is indeed more compact (lower R_g_) in complex than in the unbound state (**Fig. 3C**), which is in line with our IM-MS results (**Fig. 3B**). We determined the relative affinity of SERF for these two RNA molecules from simulations (**Fig. S4A-D**)^3,67^, which revealed a ∼3.5-fold higher affinity of SERF for TAR compared to (rU)_29_, in good agreement with the ∼2.8-fold higher affinity seen by our experiments (**Table S3)**. Although there are inherent limitations to a coarse-grained model, such simulations provide a computationally tractable way to study the sequence-specific driving forces of protein-RNA complex formation.

### Flexible regions form the SERF-TAR binding interface

To determine the nature of SERF-TAR interactions in the 1:1 complex, we again turned to NMR spectroscopy. In NMR experiments that detect on isotopically enriched TAR, we used a TAR mutant that contains a more stable tetraloop (UUCG) instead of the native hexanucleotide CUGGGA sequence^34^. For NMR analysis, the tetraloop version of TAR was used because of the complicated dynamics of the hexanucleotide loop sequence, which introduce challenges in spectroscopic interpretation^68^. The uniform ^13^C/^15^N-labeling scheme for TAR allowed us to probe the chemical environment of nuclei in both sugar and nucleobase components. From a battery of 2D TROSY-HSQC NMR experiments, we observed chemical shift differences in and around the bulge region of TAR upon the addition of SERF. For comparison, we quantified the contact frequency for each TAR bead in complex with SERF from CG-simulations and observed a qualitatively similar binding footprint at the TAR bulge (**Fig. 4A-B, S6-7**). Aside from fluctuations in the G28:C37 base pair of the upper helix peaks for the canonical Watson-Crick base pairs of the TAR helices are maintained throughout the titration. Hence, SERF binding does not appear to alter the TAR secondary structure.

**Figure 4:**
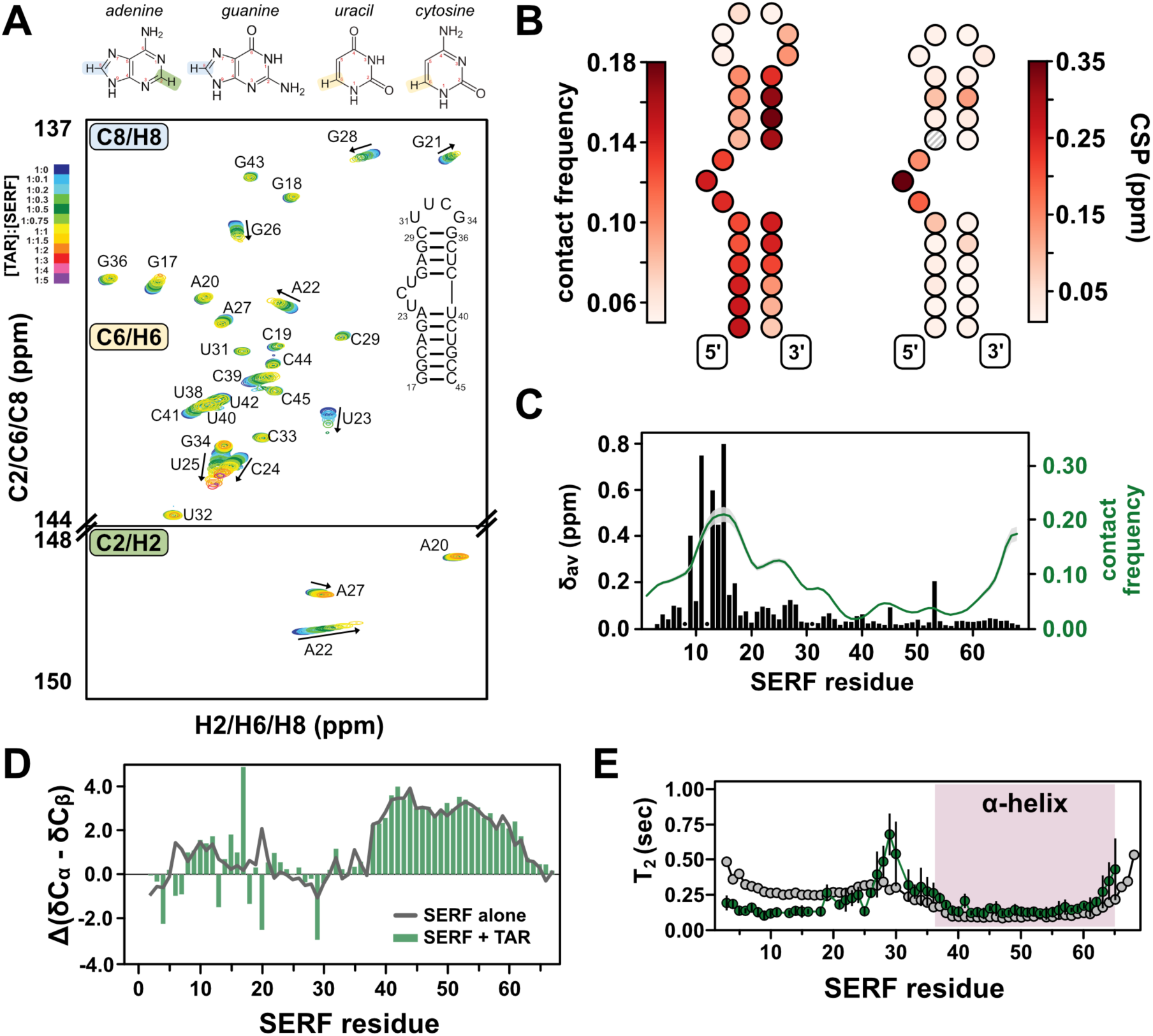
Determinants of SERF-TAR interactions. **(A)** TROSY-HSQC spectra of isotopically enriched TAR showing chemical shift changes upon titrating with unlabeled SERF (molar ratio color coded according to the insert scale). **(B)** Diagram of the binding footprint of SERF on TAR from coarse-grained simulations (left) and NMR experiments (right). Each TAR nucleotide is depicted with a circle colored to reflect either contact frequency (from simulations) or the maximal ^13^C chemical shift perturbation (CSP) (from NMR). The intensity of the coloring represents the propensity for SERF interaction at each TAR nucleotide. **(C)** Plot depicting regions of SERF that interact with TAR. Chemical shift perturbations in SERF upon adding TAR are shown as black bars (left axis). Residues for which bound-state assignments could not be determined are denoted with black dots. The average per-residue contact frequency for SERF with 15 TAR conformers is shown as a green trace; the standard deviation is shown as grey shading above and below the green line. **(D)** Secondary structure profile for TAR-bound SERF calculated from NMR chemical shifts. The unbound SERF secondary structure profile is shown as a grey line for comparison. **(E)** ^15^N transverse relaxation times for SERF in complex with TAR (green). Values for unbound SERF (gray) are reproduced from Figure 2B for comparison.

In the absence of major structural changes, the chemical shift changes in the TAR bulge region suggest that this is the likely site of SERF interaction. In support of this SERF-TAR interaction mode, other TAR binding partners, including the HIV Tat protein, also associate with the bulge region^34,69,70^. Residual dipolar coupling experiments with and without SERF illuminated changes in the relative orientation of the two TAR helices arising from SERF interaction. Consistent with observations for other TAR-protein complexes, SERF binding decreases the TAR interhelical angle from ∼52° (unbound) to ∼18° (bound) (**Table S5**)^64,71^.

We next sought to identify the region(s) of SERF that supports its interaction with RNA. To do so, we added TAR to isotopically enriched SERF and compared the spectral features of SERF with or without RNA. We observed significant line broadening in the ^13^C, ^15^N-CON spectrum of SERF in complex, which posed a barrier to resonance assignments and interpretation of subsequent experiments (**Fig. S6A-B**). Therefore, we used ^1^H, ^15^N-HSQC experiments to monitor TAR-induced chemical shift differences in the SERF backbone. We unambiguously assigned cross peaks for 62 out of the 67 non-proline residues of SERF aided by the 2D ^13^C, ^15^N-CON and 3D HNCO/HN(CA)CO spectra. The per-residue chemical shift perturbations (CSPs) arising from the addition of TAR are primarily localized to the conserved NTR (**Fig. 4C**). Recall from PRE experiments and all-atom simulations (above) that unbound SERF appears to be locally compact between residues ∼5-25. Hence, CSPs within this region collectively reflect a local de-compaction event *and* the RNA binding event.

Interestingly, despite a high density of positive residues, the lack of chemical shift perturbations in CTR suggests that this region does not interact with TAR. In support of this result, CG simulations of ‘SERF’ and ‘TAR’ show the highest contact frequency within NTR (**Fig. 4C, green trace**). (rU)_30_ binding to SERF yields systematically smaller CSPs compared to TAR binding but the perturbations are still confined to the NTR (**Fig. S6C-D**). Comparison of the CSP profiles between RNA binding partners inform a possible model in which SERF may bind with or without NTR de-compaction. Secondary structure calculations from the bound-state NMR measurements show that such features are largely unchanged from the unbound SERF ensemble (**Fig. 4D**), suggesting that the helical character in the CTR is maintained in the bound state. Taken together, the totality of data demonstrate that SERF shows no acquisition of structure upon binding to TAR. CG simulations of ‘SERF (1-34)’ with ‘TAR’ yielded the same relative dissociation constant as the full-length ‘SERF’ with TAR’ (**Table S3**), which supports the conclusion that the NTR is the primary interaction motif for these RNA partners.

We next measured transverse ^15^N relaxation times to assess the dynamics of SERF in complex with TAR. A comparison of bound versus unbound SERF relaxation profiles shows reduced backbone motions in SERF upon binding to TAR **(Fig. S4G, 4E)**. The depressed *T_2_* values of NTR in the TAR-bound versus unbound condition may reflect slower conformational exchange in this region due to its interaction with the RNA **(Fig. 1E, 4E).** Interestingly, the *T_2_* values for CTR are almost identical in the free and TAR-bound forms, which supports the conclusion that the CTR is still helical in the bound state and does not interact with RNA **(Fig. 4E)**.

Heteronuclear NOE measurements of unbound and bound SERF also highlight the differences in flexibility between N- and CTR that arise from backbone structuring and/or RNA binding (**Fig. S5E**). From the suite of NMR measurements of TAR or SERF in complex, we propose a model in which key positive residues in the NTR (i.e., Arg and Lys within the first ∼25 residues of the sequence) of SERF associate with the TAR bulge in a manner like that of other protein binding partners of TAR.

### SERF and TAR exhibit two-phase behavior at high concentrations

Given the reported localization of SERF to the nucleolus, an RNA-rich membraneless compartment, we tested whether SERF could interact with TAR to form condensates *in vitro* ^30^. Indeed, Cy3-TAR and Cy5-SERF co-localize in solution into spherical, liquid-like droplets that can wet a microscope slide surface and can fuse and then relax into a larger droplet (**Fig. 5A, S8A**). Although we observed droplet formation upon mixing SERF and TAR without a crowding agent (**Fig. S8B**), the concentration required for phase separation of SERF-TAR mixtures are about ten times lower in the presence of 10% (w/v) Polyethylene Glycol-8000 (PEG) (**Fig. 5E**). Under our experimental conditions, neither SERF nor TAR exhibited detectable phase separation behavior individually. Charge-charge repulsion, low hydrophobicity, and low aromatic amino acid content likely preclude homotypic interactions of SERF. Importantly, FITC-labeled PEG (average MW = 10K) is excluded from the SERF-TAR droplets, which confirms that PEG is not co-phase separating with SERF and TAR (**Fig. 5A**)^72^.

**Figure 5:**
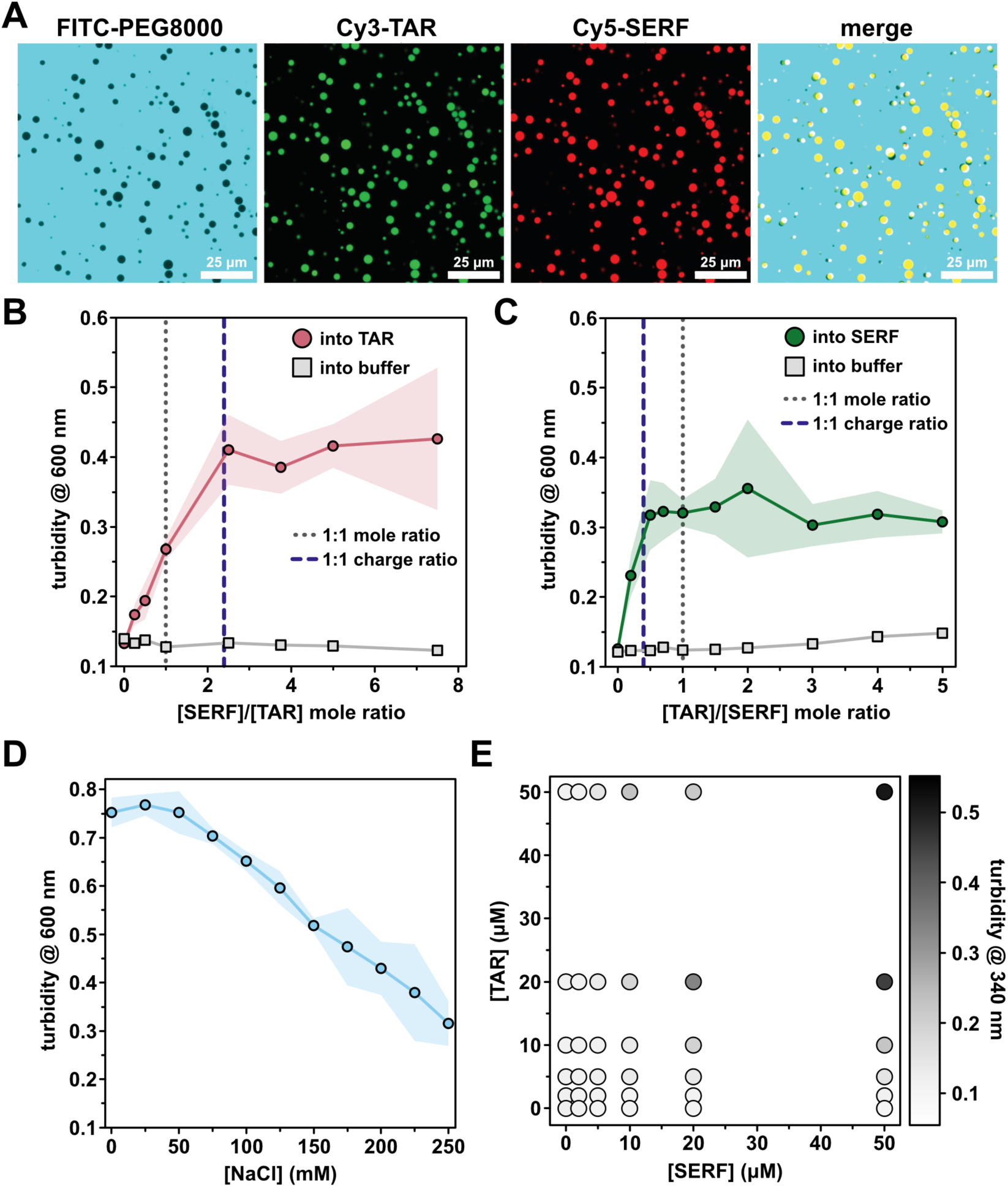
Characterization of SERF-TAR coacervates. **(A)** Fluorescence images of ySERF-Cy5 (A63C) and/or TAR-Cy3 in droplets. Samples for imaging contained 50 μM SERF and 50 μM TAR and 10% (w/v) PEG8000 in a buffer of 20 mM HEPES, pH 7.5, 85 mM NaCl, 1 mM MgCl_2_. FITC-PEG is not incorporated to droplets containing SERF and TAR. **(B)** and **(C)** Turbidity titrations **(B)** with increasing [SERF] titrated into a fixed concentration (20 μM) of TAR (pink circles) or buffer control (grey squares). **(C)** with increasing [TAR] titrated into a fixed concentration (50 μM) of SERF (green circles) or buffer control (grey squares). **(D)** Turbidity as a function of NaCl concentration. In **B**, **C**, and **D**, the shading represents the standard deviation from the average of three technical replicates. Lines connecting the data points are shown to guide the eye. **(E)** Phase diagram of SERF-TAR phase separation generated by measuring turbidity at 340 nm.

We next investigated how the ratio of SERF to TAR influenced phase behavior. We used optical turbidity measurements (λ = 340 nm) to monitor phase separation of the SERF-TAR system at various molar ratios. Increasing SERF concentrations at a fixed concentration of TAR led to saturable increase in turbidity (**Fig. 5B**). We observed similar behavior in the reverse experiment, in which TAR was added to a fixed concentration of SERF (**Fig. 5C**). At pH 7.5, SERF has an estimated charge of +12, whereas the TAR net charge is estimated to be -29. The turbidity peaked at a stoichiometric ratio of approximately 2.5:1 for SERF:TAR, which is close to the charge-matched condition (**Fig. 5B**). Thus, in the SERF-TAR system, turbidity of the sample (indicating droplet formation) begins to plateau once a 1:1 charge ratio is achieved (**Fig. 5B-C**) ^23^. In support of an electrostatically driven assembly mode, the phase separation of SERF and TAR is sensitive to ionic strength, as evidenced by a salt-dependent decrease in turbidity (**Fig. 5D, S8D**). The charge complementarity between molecules and the salt-dependence of their phase separation are hallmarks of complex coacervation, a type of associative phase separation that is driven by electrostatic attraction between oppositely charged polymers^73^. The salt-sensitive co-assembly and phase separation of SERF and TAR may be described by electrostatic driving forces alone; however, our observations do not eliminate the likely possibility that other interactions (i.e., cation-π) contribute to SERF-TAR phase behavior.

With a detailed description of the stoichiometric SERF-TAR complex and evidence for two-phase behavior at high concentrations, we sought more information about SERF-TAR assembly that precedes the emergence of a dense phase. To qualitatively assess the association of SERF monomers into higher-order species, we performed crosslinking experiments with a heterofunctional amine-to-carboxylic acid crosslinker, 4-(4,6-Dimethoxy-1,3,5-triazin-2-yl)-4-methylmorpholinium (DMTMM). DMTMM is a zero-length crosslinker that can couple acidic residues (Asp, Glu) with primary amines (usually Lys)^74^. In the absence of RNA, SERF migrates as a monomer on a denaturing SDS-PAGE gel. This result is expected, given the charge and sequence features of SERF, and consistent with our biophysical measurements at relatively high SERF concentrations that do not implicate homotypic SERF assembly (i.e., NMR, SAXS). In crosslinking experiments containing SERF and TAR, we observed crosslinked, higher-order SERF species that migrate distinctly from the monomer (**Fig. S8C**). The smallest species in mixtures containing RNA is monomeric SERF. The higher molecular weight bands presumably represent crosslinked SERF dimers and other oligomeric species (**Fig S8C**). Although crosslinking approaches cannot distinguish between direct intermolecular contacts and spatial proximity of reactive groups on SERF in the presence of RNA, these results suggest that soluble SERF-TAR may explore higher-order assemblies in the background of a primarily binary complex.

## DISCUSSION

In this study, we have investigated the *in vitro* interactions of a small, disordered protein, SERF, and a fragment of the HIV-1 TAR RNA and established this pair as a tractable model system for studying IDR interactions with RNA. SERF lacks a canonical, folded RNA-binding domain and instead leverages its high fraction of positive amino acids toward RNA binding.

SERF also contains a transient α-helix in its CTR, but this region has no known functions or binding partners and does not appear to influence or be influenced by RNA binding. At high concentrations of SERF and RNA (or with the addition of a crowding agent), the pair undergo complex coacervation to form a protein- and RNA-rich dense phase. Importantly, the threshold for SERF-TAR phase separation is sufficiently high that we could use NMR spectroscopy to study the 1:1 complex to gain residue-level insights into this interaction. NMR measurements of SERF or TAR in complex suggest that neither molecule changes in secondary structure upon binding. Consistent with studies of TAR with other binding partners, we found that binding to SERF induces re-orientation of stem loop helices around the TAR bulge^64^. From IM-MS experiments and coarse-grained simulations, the SERF ensemble appears to modestly compact upon binding to RNA.

We find no evidence for higher-order assembly of SERF in the absence of RNA, which is likely a consequence of its charge and sequence features that make self-assembly electrostatically disfavored. From polymer theory, complex coacervation of oppositely charged polymers nominally consists of two thermodynamic steps: (1) entropically favored pairing between the positively and negatively charged polymers followed by (2) higher-order assembly into complex coacervates^75^. Introducing a polyanion (RNA) effectively neutralizes the positive charges that preclude SERF-SERF interactions. This is captured by IM-MS experiments and CG simulations as a modest compaction of the SERF ensemble in the RNA-bound state. Mitigation of *intra*molecular repulsion by RNA binding should extend similarly to *inter*molecular repulsion between SERF molecules. This may ultimately support the emergence of SERF-SERF contacts that enhance the multivalency required for phase separation.

The electrostatic interactions and dynamics underlying complex formation and phase separation of two highly and oppositely charged disordered proteins was recently interrogated for the histone chaperone Prothymosin α and histone H1^76,77^. At low concentrations, the pair form an extremely high-affinity (∼ picomolar) binary complex. With increasing concentrations, Prothymosin α and histone H1 undergo liquid-liquid phase separation and form salt-sensitive, viscous droplets that are stabilized by the same electrostatic interactions as those which stabilize the 1:1 complex^77^. A recent investigation into RBP-RNA interactions within biomolecular condensates revealed that the sequence-specific RBP-RNA contacts that support dilute-phase interaction are also present in the dense phase^78^. This study also reported the emergence of additional protein-RNA contacts in the dense phase arising, in part, from non-specific, electrostatic interactions of RBP IDRs with the RNA phosphate backbone.

Whether this paradigm holds true for IDR-RNA interactions generally is unknown, but our finding that SERF compacts upon binding to RNA supports such a model for emergent contacts in higher-order assemblies via attenuated electrostatic repulsion. We note similarities between our system and a disordered protein that facilitates folding of its nucleic acid binding partner^4^. This study reported accelerated folding kinetics of a DNA hairpin via non-specific, electrostatic interactions with a highly positively charged disordered protein. In this system, the disordered protein acts as the counterion to permit nucleic acid compaction and base-pairing.

Supporting the idea of charge screening as the primary factor in compaction, high-salt conditions recapitulated the compaction and increased rate of DNA folding^4^. Although SERF does not fold upon binding RNA, the modest compaction of the ensemble implicates the same charge screening effect that promotes DNA folding upon binding to the polycationic protein.

For many RBPs, folded domains and intrinsically disordered regions jointly contribute to RNA-binding and phase-separation abilities^3,66,79^. However, a consequence of high multivalency is that many such proteins readily phase separate, which poses a substantial barrier to characterize the residue-level details of the interactions^12,26,80^. Proteome-wide studies to identify RBPs uncovered a remarkable fraction of binding sites that map to IDRs^81,82^. Thus, understanding the modes of interaction between IDRs and RNA will inform how RBPs with and without folded RNA-binding modules leverage IDRs in RNA recognition and binding. Through small IDR-RNA systems, like SERF-TAR, we may gain insights into the residue-level interactions that support a dilute-phase complex.

Recent studies of biomolecular condensate nucleation and early-stage assembly revealed that phase-separating proteins can form small ‘clusters’ in the dilute phase that vary in the number of constituent molecules^83–85^. The path from monomers to clusters to condensates is energetically complex, but the formation of relatively low-affinity clusters seems to be the first kinetic and energetic barrier to subsequent higher-order assembly^85^. Although this model emerged from protein-only studies, we see elements of a similar assembly path in crosslinking experiments of SERF-TAR by the RNA-dependent formation of higher-order SERF species in the dilute phase. It is our hope that SERF and TAR (or unstructured r(U)_30_) may be a tractable system to enable further exploration of the general features of IDR-RNA interactions that support the various stages of condensate assembly.

## AUTHOR CONTRIBUTIONS

R.M., E.T.U., S.D., N.H.Y, V.V.G. and J.C.A.B designed the research and analyzed the data. R.M., S.D., E.T.U., M.J.C., N.H.Y., O.A.F., J.G., V.V.G. performed the experiments. S.A.S., A.S.H., B.T.R., L.S., and J.C.A.B. supervised the research. R.M., E.T.U., S.D., N. H.Y., and J.C.A.B. wrote the manuscript. All authors edited the paper.

## DATA AND CODE AVAILABILITY

The source data for the figures and supplementary figures are included as a Source data file and also deposited in Open Source Framework (https://osf.io/6qzhk/). All other data are available from the corresponding author upon reasonable request.

## MATERIALS AND METHODS

### Oligonucleotides

Unless otherwise specified, the HIV-1 TAR RNA used for experiments has the sequence 5′-GGCAGAUCUGAGCCUGGGAGCUCUCUGCC-3′^86^. Unlabeled TAR RNA from Horizon Discovery and Cy3 3′-labeled TAR RNA from Integrated DNA Technologies were purchased as HPLC-purified lyophilized solids and were used without further purification. All solutions were prepared in nuclease-free water (Ambion) and buffers were prepared in DEPC-treated double-distilled water to remove RNases.

Typically, labeled RNA was dissolved in nuclease-free water to a stock concentration of 100 µM. The RNA was annealed by heating at 95 °C for 5 min followed by rapid cooling by plunging into ice. U30 RNA (Integrated DNA Technologies) was resuspended in 50 mM potassium phosphate (pH 6.5), 50 mM KCl, 1 mM MgCl2, 0.01% NaN3 buffer and used directly (i.e., no annealing protocol) for NMR experiments.

### 13C-15N labeled TAR RNA preparation

The HIV-1 TAR containing a UUCG loop instead of the CUGGGA loop was produced by *in vitro* transcription with T7 RNA Polymerase High-Concentration (New England Biolabs) in 40 mM Tris HCl (pH 8), 1 mM Spermidine, 0.01% TritonX-100, 5 mM DTT, 28 mM MgCl2, and 4mM ^13^C,^15^N-enriched NTPs (Eurisotop). The transcription reaction ran for 4 hours at 37°C on a heater/mixer block (Eppendorf ThermoMixer), in the presence of 240 nM of semi-double stranded DNA template containing two 5’ C2’-methoxy nucleotides. The reaction was quenched with 10% v/v 0.5 M EDTA. The purification was performed by ion-exchange high pressure liquid chromatography (HPLC) (Agilent) and using DNA-Pac PA1000 column at 80 °C (Thermo Scientific). The elution was performed by using a buffer containing 12.5 mM Tris (pH 8.0), 8 M urea, and a gradient of NaCl (from 0 to 0.5 mM) and detected by UV signal at 260 nm. Fractions were pooled together, butanol concentrated, and then precipitated by ethanol and sodium acetate overnight. The sample was washed with ethanol and remaining traces of ethanol were evaporated using a SpeedVac. The sample was resuspended in NMR Buffer (15 mM potassium phosphate buffer (pH 6.4), 50 mM NaCl, 0.1 mM EDTA, 10% D2O), denatured by heating at 95 °C for 5 minutes, and cooled down at room temperature. The concentrations of RNA samples were measured by using NanoDrop 2000 spectrophotometer and adjusted to a final concentration of 0.5 mM.

### Protein expression and purification

The *Saccharomyces cerevisiae YDL085C-A* gene that encodes the SERF protein was codon-optimized for expression in *E. coli*. Point mutations were inserted into the SERF gene by site-directed mutagenesis with a QuikChange kit (Agilent) and were verified by Sanger DNA sequencing. The wildtype and mutant SERF genes were cloned into a pET28 vector that encodes an N-terminal His6-SUMO fusion tag for purification and scarless cleavage by ULP1 protease^63^. The plasmid was transformed into *E. coli* BL21 (DE3) cells for protein expression. Cells were grown to early logarithmic phase at 37 °C in Protein Expression Medium (Gibco) or M9 minimal medium supplemented with ^15^NH4Cl (for expressing uniformly ^15^N labeled protein), both containing 100 µg/mL kanamycin. Prior to induction, cells were cooled to 20 °C and then expression was induced by adding 0.1 mM isopropyl β-D-1-thiogalactopyranoside (IPTG). After 16 hours of protein expression at 20 °C, cells were harvested by centrifugation and resuspended in lysis buffer containing 40 mM Tris, 10 mM sodium phosphate (pH 8.0), 10% glycerol, three tablets of protease inhibitor cocktail (cOmplete mini EDTA-free, Roche), 0.375 mM MgCl2, and 0.05 μg/ml each DNase I and RNase A. Cell pellets containing SERF were lysed by sonication on ice for 8 min, followed by lysate clarification via centrifugation twice at 37,500 ×g for 30 min at 4 °C. The clarified supernatant was loaded on a 5 mL HisTrap prepacked column (Cytiva) that had been equilibrated with lysis buffer. The column was washed with Lysis Buffer and the His6-SUMO tagged protein was eluted by adding Lysis Buffer containing 500 mM imidazole. To remove any undegraded nucleic acids, 500 units of Pierce™ Universal Nuclease (Cat# 88701) and 1 mM MgCl2 were added to the eluate. ULP1 and 10 µL β-mercaptoethanol were also added to the eluted material to initiate SUMO tag removal. The resulting mixture was dialyzed against 40 mM Tris (pH 8.0), 300 mM NaCl overnight at 4 °C. The cleaved affinity tag was separated from the SERF by passing the protein solution over a HisTrap column that had been equilibrated with Lysis Buffer. The flow-through fraction from the column containing the tag-free SERF was diluted by adding a cation exchange buffer (50 mM sodium phosphate (pH 6.0), 100 mM NaCl). Following centrifugation to remove insoluble material, the protein was passed over a HiTrap SP cation exchange column (Cytiva) that had been equilibrated with the cation exchange buffer. The protein was eluted with a linear gradient using a buffer containing 1 M NaCl. The eluted fractions were concentrated and loaded on a HiLoad Superdex S75 gel filtration column (Cytiva) equilibrated with 40 mM HEPES (pH 7.5), 100 mM NaCl. Purified SERF was flash-frozen in liquid nitrogen, and either aliquoted and stored at -80 °C, or subjected to overnight dialysis in 50 mM ammonium bicarbonate pH 8.0 at 4 °C followed by freeze-drying and storage at -80 °C. Protein concentration was determined with the Qubit protein assay (ThermoFisher Scientific) following manufacturer’s protocol. Uniformly ^15^N and ^13^C isotope-enriched proteins were expressed following the protocol of Marley et al. in M9 minimal medium supplemented with ^15^NH4Cl, U-^13^C-glucose, and ISOGRO growth supplement (Sigma)^87^.

### Fluorescence anisotropy

The apparent dissociation constant for the SERF-TAR complex was determined by titrating SERF (0.5 mM stock) into a solution containing 15 mM sodium phosphate (pH 6.4), 50 mM NaCl, 0.1 mM EDTA, and 200 nM 3’6-FAM-labeled TAR RNA (100 µM stock) that was heated to 95 °C for 5 min and then cooled rapidly in an ice bath for 10 min or until use. Fluorescence signal was recorded at 25 °C with a Cary Eclipse Spectrofluorometer (Agilent) using excitation and emission wavelengths of 493 nm and 520 nm (5 nm bandpass for both) respectively, and anisotropy values were calculated using the following equations:

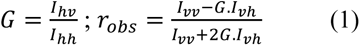

where *G* is an instrument correction factor, *I* is the measured fluorescence intensity with polarizers oriented in directions that are indicated in subscripts (*v* is vertical and *h* is horizontal) and *robs* is the measured anisotropy. The titration points could be fitted to a Langmuir-type isotherm describing the fraction of ligand (SERF) that is bound (FB):

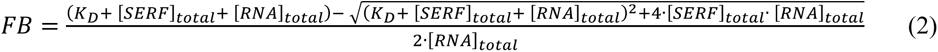

where [SERF]total and [RNA]total are the concentrations of the ligand (SERF) and macromolecule (RNA), respectively, and KD is the fitted dissociation constant (in the same units as the macromolecule and ligand concentrations). Fraction bound from Eq. 2 relates to the spectroscopic observable (*robs*) through normalization based on the range of the measured anisotropy values:

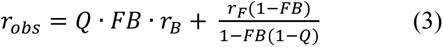

Where *rF* is the anisotropy of the free fluorescent molecule, *rB* is the anisotropy of the bound fluorescent molecule, and *Q* is a dimensionless correction value to account for differences in absolute fluorescence intensities between free and bound states. Using the measured *robs,* the fluorescent macromolecule (RNA) concentration (constant), and the total concentration of ligand (SERF) for each data point, we extracted KD and *Q* by non-linear least squares fitting in Python.

### Spin-Labeling Cysteine Mutants

Because wild-type SERF lacks native cysteines, we expressed and purified variants containing single cysteines at position 63 (A63C) or 10 (A10C) for attachment of the paramagnetic nitroxide spin label, MTSSL (1-Oxyl-2,2,5,5-tetramethylpyrroline-3-methyl methanethiosulfonate). MTSSL stock was prepared in acetone at a concentration of 100 mg/mL. Uniformly ^13^C, ^15^N -SERF A10C sample was diluted to 1 mL in Tris buffer, and freshly prepared DTT was added to a final concentration of 10 mM (>10-fold molar excess). The SERF samples were incubated for at least 1 hour at 4 °C in the dark to reduce cysteine. The samples were then buffer-exchanged into 50 mM potassium phosphate (pH 6.5), 50 mM KCl, 1 mM MgCl2, 0.01% NaN3 using a PD-10 buffer exchange resin (Cytiva) to remove the excess DTT. MTSSL was added to a final concentration of 1 mg/mL, and the sample was incubated for 1 hour at room temperature (∼22°C) in the dark. The unconjugated MTSSL was removed from MTSSL-labeled SERF samples using a PD-10 resin equilibrated with 50 mM potassium phosphate (pH 6.5), 50 mM KCl, 1 mM MgCl2, 0.01% NaN3. Labeling was confirmed by mass spectrometry.

### NMR spectroscopy

The acquisition parameters for experiments involving SERF chemical shift assignment, spin relaxation measurements, and PRE experiments are listed in **Table S6**.

#### Chemical Shift Assignment and Secondary Structure Analysis

Uniformly labeled ^13^C, ^15^N -SERF was buffer-exchanged into 50 mM potassium phosphate (pH 6.5), 50 mM KCl, 1 mM MgCl2, 0.01% NaN3 using a desalting column and concentrated to 0.8 -1 mM. Samples for NMR experiments were made to a volume of 500 µL with 5 % D2O for the deuterium lock. All NMR spectra were collected at 27 °C on a Bruker Avance NEO 600 (^1^H) MHz spectrometer with a 5 mm TCI triple-resonance cryoprobe. On-instrument processing of all spectra was performed using the Bruker Topspin software. Peak picking and manual assignments were carried out in NMRFAM-Sparky^88^.

Resonance assignments were acquired through ^13^C direct-detect spectra^37^. Nearest-neighbor correlations between amide nitrogen atoms on adjacent residues were established using 3D (HACA)N(CA)CON and (HACA)N(CA)NCO experiments and 2D amino acid-filtered experiments collected as CACON/CANCO spectra for Asp, Ala, Glu, and CACON spectrum for Leu/Ala resonances^37^. Side-chain chemical shifts were assigned by collecting 3D CCCON spectra for aliphatic carbon and 3D H(CC)CON aliphatic proton resonances^37^. The resonance assignments were mapped onto the ^1^H,^15^N-HSQC spectrum using standard 3D HNCACB, CBCA(CO)NH, HNCO, and HN(CA)CO spectra. Secondary structure propensity was determined by calculating residue-specific secondary chemical shifts, i.e., the difference in the measured ^13^C^α^ and ^13^C^β^ chemical shifts from the sequence-specific random coil chemical shifts calculated by the *ncIDP* predictor for the SERF amino acid sequence^89^.

#### Spin Relaxation Measurements

^15^N *T1* and *T2* spin relaxation data were collected as pseudo-3D spectra using ^1^HN-start CON experiments developed previously for ^13^C direct-detect spin relaxation measurements^45^. For *T1* measurements, 13 fully interleaved spectra were collected with relaxation delays of 20, 50, 80, 100, 150, 180, 200, 300, 400, 600, 750, 800, 1000, and 2000 ms. For *T2* measurements, 16 fully interleaved spectra were collected with relaxation delays of 15.68, 31.36, 62.72, 78.4, 94.08, 109.76, 125.44, 156.8, 172.48, 188.16, 219.52, 235.2, 250.88, 282.24, and 313.6 ms (the asterisks indicate duplicate measurements). The decay curves were fitted to a single exponential function (**Eq. 4**). Reported error bars represent uncertainty in fitting for each data point.

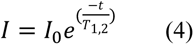

#### Paramagnetic Relaxation Enhancement (PRE) Measurements

A ^1^H-start variation of the CON spectrum ((HACA)CON) was acquired on the freshly prepared paramagnetic sample containing spin-labeled A10C or A63C SERF in 50 mM potassium phosphate (pH 6.5), 50 mM KCl, 1 mM MgCl2, 0.01% NaN3 supplemented with 5% D2O. Following data acquisition, the MTSSL radical was quenched by mixing 1 µL of 0.5 M sodium ascorbate directly into the NMR tube containing 500 µL of the sample. Following incubation at room temperature for 10 minutes, identical NMR spectra were acquired for the diamagnetic sample. PREs were reported as the ratio of cross-peak intensities under paramagnetic (*Ipara*) and diamagnetic conditions (*Idia*).

#### Protein NMR experiments on the SERF-TAR complex

The NMR samples contained 100 µM ^13^C, ^15^N-SERF and 100 µM TAR or 100 µM U30 in 50 mM potassium phosphate (pH 6.5), 50 mM KCl, 1 mM MgCl2, 0.01% NaN3 buffer supplemented with 8% D2O. Resonance assignments of SERF in the presence of TAR were generated based on the traditional ^1^H, ^15^N HSQC experiment. The 3D experiments that were collected for assignments include HNCO, HN(CA)CO, CBCA(CO)NH, and HNCACB. Spectral comparison was carried out using the assigned HSQC spectrum of SERF alone as a reference. ^15^N relaxation experiments on SERF in complex with RNA were conducted using 100 µM ^15^N-SERF and 100 µM TAR. The *T2* measurement was carried out with the same relaxation delays as for SERF alone. The heteronuclear two-dimensional {^1^H} −^15^N nuclear Overhauser effect (hetNOE) was measured with a relaxation delay set to 5 seconds. The hetNOE profile was determined from the ratio of peak heights in the experiments with and without proton saturation.

#### SERF titrations and RDC measurements of TAR RNA

All RNA NMR experiments were acquired with 0.5 mM ^13^C, ^15^N-enriched TAR and varying concentrations of SERF in 15 mM Phosphate buffer (pH 6.4), 50 mM NaCl, 0.1 mM EDTA, and 10% D2O at 25°C on Bruker spectrometer equipped with a QCI cryoprobe and operating at 600 MHz ^1^H frequency. The titrations of SERF were observed through 2D ^15^N- or ^13^C-TROSY-HSQC spectra at concentrations of 0.05, 0.1, 0.15, 0.25, 0.375, 0.5, 0.75, 1, 1.5, 2, and 2.5 mM SERF. This range of concentrations resulted in TAR:SERF mole ratios spanning from 1:0.1 to 1:5. Combined chemical shift perturbation (CSP) (Δδ), were computed by:

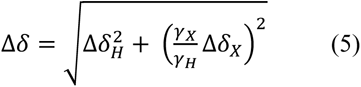

where *γ*_H_ and *γ*_X_ are the gyromagnetic ratio of ^1^H and X (either ^13^C or ^15^N) and ΔδH and ΔδX the changes in chemical shift measured for the given nuclei (in ppm). Only peaks with a signal to noise ratio above 5 were kept in the analysis. Maximal ^13^C CSPs correspond for each nucleotide to the maximal CSP observed for C1’H1’, C2H2, C5H5, C6H6 or C8H8 in the 1:1 complex or to the latest observable fraction if the signal was unobservable at 1:1. The apparent dissociation constant of the interaction assuming a 1:1 complex was determined by non-linear fitting.

TAR RDCs (N1H1, N3H3, C1’H1’, C2H2, C5H5, C6H6 and C8H8) were measured using pairs of 2D TROSY semi-anti-TROSY HSQC spectra recorded on a 1:1 TAR:SERF complex in presence of 25 mg/ml Phages (ASLA), obtained by mixing a 1 mM 1:1 complex with an equal amount of Phages (50 mg/ml in 15 mM Phosphate buffer pH 7.4). All spectra were processed using NMRPipe and analyzed in CcpNMR^90,91^.

RDC data were analyzed using Module ^90–92^ assuming idealized A-form helices for the two helices that were analyzed independently. Scalar products between alignment tensors were computed as previously described ^64^. It is worth noting that the RDCs were recorded on a 1:1 complex, which appear not to be saturated under NMR conditions and within a fast-intermediate exchange regime. Consequently, the obtained values do not report on the complex, but on a population weighted average of the free and bound form ^93^. Therefore, only qualitative trends were extracted from this analysis. RDCs for the free TAR were taken from the literature^64^.

### Preparation of SERF-RNA condensates

SERF was reconstituted from lyophilized powder into the LLPS buffer (20 mM HEPES/NaOH (pH 7.5), 85 mM NaCl, 1 mM MgCl2). SERF was reconstituted from lyophilized powder into the LLPS buffer (20 mM HEPES/NaOH (pH 7.5), 85 mM NaCl, 1 mM MgCl2). TAR RNA was reconstituted in LLPS buffer and concentration was measured by absorbance at 260 nm in a Nanodrop ND-1000 spectrophotometer. Labeled TAR was reconstituted in RNase-free water and diluted into working solutions to the desired concentration. Protein concentrations were measured using Invitrogen Qubit Protein Assay kit IAW on a Qubit 2.0 fluorometer. Samples were prepared by sequentially combining LLPS buffer with 10 % PEG 8000 (w/v), TAR, and then SERF. Upon addition of SERF, 20% of sample volume was pipetted slowly for adequate mixing. All components were maintained at room temperature during sample preparation. For conditions in which phase separation was observed, sample turbidity increased concomitant with mixing indicating the rapid formation of two phases.

#### Microscopy

Condensate samples were added into Corning 96-well microplates with glass bottom (Cat#4580) pretreated overnight with 5% (w/v) Pluronic acid and rinsed thoroughly with ddH2O. All fluorescent imaging were performed on a Leica SP8 confocal microscope equipped with LAS X Life Science Microscope Software and 100X objective lens. Phase-separated droplets contained 1 % labeled components. 3’Cy3-labeled RNA was imaged using 554 nm excitation and 568 nm emission. Cy5 labeled SERF A63C mutant was imaged using 651 nm excitation and 670 nm emission. mPEG-FITC, MW 10 K (Creative PEGWorks Cat# PSB-2253-100 mg) was imaged using 495 nm excitation and 519 nm emission.

#### In vitro crosslinking

50 mM 4-(4,6-Dimethoxy-1,3,5-triazin-2-yl)-4-methylmorpholinium Chloride (DMTMM) stock solution was freshly prepared by dissolving 13.84 mg of DMTMM powder in 540 μL of anhydrous dimethyl sulfoxide. DMTMM solution was added to a final concentration of 1.25 mM to samples containing different concentrations of SERF and TAR in LLPS buffer (20 mM HEPES/NaOH (pH 7.5), 85 mM NaCl, 1 mM MgCl2). The crosslinking reaction was allowed to proceed for 1 hour at room temperature. Excess unreacted DMTMM was quenched by addition of 1 M Tris pH 7.5 solution to a final concentration of 100 mM Tris. The quenching reaction was incubated for 45 min at room temperature. The residual crosslinker was removed by centrifugation and 100 μL supernatants were mixed with 25 μL of 5X reducing SDS-sample buffer. The samples were boiled at 95 °C for 10 min and run on a NuPAGE 4-12% Bis-Tris polyacrylamide gel (Thermo Fisher Cat#NP0322BOX). Bands were visualized by staining the gel with Pierce GelCode blue stain reagent following manufacturer’s protocol.

#### Small-angle X-ray scattering

All SAXS data were collected at 20 °C. SERF samples for SAXS were buffer-exchanged into 50 mM potassium phosphate (pH 6.5), 50 mM KCl, 1 mM MgCl2, 0.01% NaN3. Small angle X-ray scattering (BioSAXS) was collected on SERF (at 4.5 and 9 mg/ml), TAR RNA (at 3 and 5 mg/ml) and SERF-TAR complex at 250 µM. The concentration of SERF samples was determined using a Direct Detect Fourier transform infrared spectrometer (EMD Millipore). SAXS data were collected on freshly eluted samples from a Wyatt S-100 size-exclusion chromatography column using X-rays generated by a Rigaku MM007 rotating anode housed with the BioSAXS2000^nano^ Kratky camera system at a wavelength (λ) of 1.54 Å. The system includes OptiSAXS confocal max-flux optics that is designed specifically for SAXS and a sensitive HyPix-3000 Hybrid Photon Counting detector. The sample capillary-to-detector distance was 495.5 mm and was calibrated using silver behenate powder (The Gem Dugout, State College, PA). The useful momentum transfer scattering vector q-space range (*q* = 4.π.sin(θ)/λ , such that 2θ is the scattering angle) was generally from qmin= 0.008 Å^-1^ to qmax= 0.6 Å^-1^. The energy of the X-ray beam was 1.2 keV, with the Kratky block attenuation of 22% and a beam diameter of ∼100 μm. Protein samples were loaded using the Rigaku autosampler into a quartz capillary flow cell mounted on a sample stage cooled to 4°C and aligned in the X-ray beam. The sample cell and full X-ray flight path, including beam stop, were kept *in vacuo* (< 1×10^-3^ torr) to eliminate air scatter. The Rigaku SAXSLAB software was programmed for automated data collection of each protein with elaborate cleaning cycles between samples. Data reduction including image integration and normalization and background buffer data subtraction were also carried out using the SAXSLAB software. Six ten-minute images and three replicates from protein and buffer samples were collected and averaged after ensuring that no X-ray radiation damage had occurred. SAXS data overlays showed that there was no radiation decay or sample loss over the 60 minutes of data collection. This was followed by reference buffer subtraction to get the raw SAXS scattering curve from only the RNA. The data were analyzed using the ATSAS API^94^.

#### Turbidity measurements

Turbidity measurements were taken at either fixed 20 µM of TAR with varying SERF concentrations and 50 µM of SERF with varying TAR concentrations. Fifty microliters of sample were added to wells in a 384-well, transparent, non-binding microplate from Greiner Bio-One (REF: 781901). Turbidity (light scattering) at room temperature was monitored as absorbance at 340 nm or 600 nm wavelengths using a Tecan M1000 infinite microplate reader. The effect of varying ionic strength was examined by adding small volumes of 5 M NaCl prepared in DEPC treated ddH2O water to 50 µL solutions containing 50 µM of SERF and 50 µM of TAR in LLPS buffer.

#### Native IM-MS

SERF and TAR were reconstituted in 100 mM ammonium acetate (pH 7.5). TAR RNA was prepared by heating the RNA at 95°C for 5 min. After heating, magnesium acetate was spiked in at a final concentration of 2 mM, and the mixture was slowly cooled to 25°C for 30 minutes to allow the RNA to refold into a favored native-like structure. The RNA was exchanged into fresh 100 mM ammonium acetate (pH 7.5) to remove excess magnesium acetate while retaining the Mg^2+^ ions bound to the RNA. Refolded TAR and SERF were mixed at a 1:1 stoichiometric ratio, with final concentrations of 4.5 μM each. The mixture was co-incubated for 30 minutes and analyzed by native ion mobility-mass spectrometry. Buffer exchanges were performed using Micro Bio-spin 6 size exclusion spin columns (Bio-Rad, Hercules, CA). Native ion mobility-mass spectrometry measurements were performed on a modified Agilent 6560c drift tube ion mobility quadrupole time-of-flight mass spectrometer optimized for native biomolecular measurements (Agilent Technologies, Santa Clara, CA)^95–97^. The instrument was operated in positive polarity, and under 99.999% nitrogen gas. Samples were introduced via nano-Electrospray ionization using 1300 kV of capillary voltage. The instrument analysis settings were optimized to preserve native-like structure and non-covalent interactions. Briefly, the source desolvation gas temperature was operated at 20 °C, and gas flow was reduced to 1.5 L/min. The front funnel, trapping funnel, drift tube, and time-of-flight tube were operated at 4.94, 3.80, 3.95, and 1.54 x 10^-7^ torr respectively. The drift tube was operated with an entrance voltage of 1700 V and an exit voltage of 250 V, enabling a low-field condition (18.125 V/cm). The IM arrival time distributions of ions were fit to gaussian functions, and the centroids of the fit gaussian functions were converted to rotationally averaged collision cross section (^DT^CCSN2) using the previously described single-field calibration using Agilent Tune Mix ions ^98^. IM measurements were performed in nitrogen, however for ease of comparison with previously published data, the CCS measurements were converted to ^DT^CCSHe using a previously established relationship^99^. Both ^DT^CCSN2 and ^DT^CCSHe are reported in the supplemental information (**TABLE S4**). All data was analyzed using Agilent IM-MS Browser 10. Raw IM data extraction, and gaussian fitting were performed using CIUSuite2^100^.

### Coarse-grained (CG) Simulations in LAMMPS

#### LAMMPS simulations with the Mpipi forcefield

Coarse-grained molecular dynamics simulations of SERF with or without a given RNA molecule were performed using the LAMMPS simulation engine and the physics-driven Mpipi forcefield^82^. In this model, each amino acid or nucleic acid monomer is represented as a single bead. Although we refer to the molecules in these simulations as ‘protein’ and ‘RNA’, it is more accurate to consider them as “protein- and RNA-flavored polymers”. In Mpipi, lysine and arginine residues harbor relative charges of +1; aspartate, glutamate, and each nucleotide have relative charges of -1. Ionic strength is represented implicitly in Mpipi; for the simulations herein, the effective NaCl concentration was set to 0.05 M. This ionic strength allowed for the protein and RNA molecules to undergo several association and dissociation events over the course of the simulation.

Prior LAMMPS simulations of the single-stranded RNA (ssRNA) homopolymer rU40 showed excellent agreement with SAXS measurements of the same sequence^67^. Our simulations use rU30 to be consistent with *in vitro* binding experiments. Unless otherwise specified, SERF and rU30 are represented as flexible polymers and are permitted to sample physically relevant conformations. Importantly, we observe remarkable agreement in the SERF ensemble dimensions between SAXS experiments, all-atom (AA) simulations, and coarse-grained (CG) simulations of SERF alone (Rg^SAXS^ = 24.9 ± 0.1, Rg^AA^ = 25.9 ± 0.8, Rg^CG^ = 25.4 ± 0.1).

#### Simulations of rU30 ssRNA with SERF

Simulations were performed using a 40 nm^3^ box with periodic boundaries. Unless stated otherwise, all simulation replicates were run for 400 million steps using a timestep of 10 fs. The first 0.25% (1 million) of the steps was discarded as equilibration steps and molecule coordinates were saved every 20,000 steps. Each system was simulated with ten independent replicates using the above settings; for SERF-rU30, each replicate used a different random starting conformation to generate final trajectories of >199,000 total frames.

#### Simulations of CG ‘TAR’ RNA with SERF

Simulations containing structured TAR molecules were performed with the same simulation parameters but used a rigid CG representation of the TAR RNA instead of a flexible polymer. The Mpipi force field was initially developed to model biomolecular liquid-liquid phase separation and so is especially well-suited to study the SERF-RNA interactions. Given this specific utility, Mpipi was parameterized to model fully flexible polymers, not folded biomolecules. Therefore, to preserve its 3D structure, TAR was only allowed global rotational and translational moves (i.e., no sampling of conformational space at the residue/bead level). To account for this, we began with a published NMR structure of TAR (PDB ID: 1ANR) and used each of the 20 deposited conformers as a TAR structure for these simulations. Each conformer was simulated with flexible SERF in five independent replicates. Each replicate used the same rigid TAR conformer paired with a random starting conformation of SERF.

We converted the atomic coordinates of each TAR conformer PDB file into ‘coarse-grained’ format by removing atoms such that each nucleotide contained a single carbon atom (C1’). This is analogous to the use of Cα coordinates to generate a coarse-grain model of a folded protein domain from a PDB file^3,67^. It is important to acknowledge that, due to how nucleotides are represented in Mpipi, there is little difference between A, U, C, and G beads. And considering the previously demonstrated agreement in the behavior of rU40 in Mpipi and experiments, we opted to use only rU beads in the rigid representation of TAR^67^. This choice is supported by the remarkable consistency of SERF-TAR simulations with expectations based on SERF-rU30 simulations and SERF-TAR experimental measurements (*discussed in next section*).

### CG Simulation Processing and Analysis

#### Calculation of relative dissociation constants from simulations of 1:1 SERF: RNA complexes

Using the simulation parameters described above, we observed numerous binding and unbinding events between SERF and rU30 over the course of the simulation. The method used for determining an apparent KD from CG protein-RNA simulations has been reported previously ^3^. We first constructed a ‘center-of-mass (COM) trajectory’ in which each molecule is represented as a single bead whose position is given by the center of mass of the CG molecule at each frame. From each COM trajectory, we constructed a radial distribution function representing the intermolecular distances across all frames. The bimodal nature of this distribution arises from distance measurements between either two bound (short distance) or two unbound (long distance) molecules. The distribution was fitted with a two-Gaussian model. The threshold distance that defines whether the complex is ‘bound’ or ‘unbound’ was determined based on the intersection of the two Gaussians. Although the threshold distance nominally depends on the molecule sizes, the use of the same protein sequence and similar RNA lengths in different simulations allows us to assume the same threshold distance across our analyses here. A threshold distance of 53.5 Å (determined from the SERF-rU30 simulations) was used for all conformations of the SERF-TAR system. Frames were defined as ‘bound’ if the COM-COM distance was < 53.5 Å (**Fig. S4A**) and five or more consecutive frames fell below this threshold. The persistence of a complex for five or more frames was used as a proxy for the ‘lifetime’ that sets a true complex apart from a stochastic or random intermolecular encounter.

The fraction of assigned bound frames was used to calculate an apparent dissociation constant with analogy to the calculations from using the second Virial coefficient to account for the finite size of the simulation box^101–103^. The analyses presented from the CG simulations of SERF and TAR reflect the averages over 15 (of the original 20) conformers. To ensure the most rigorous analyses, we omitted simulations of TAR conformers that do not adequately sample the unbound state (**Fig. S4D**). All SERF-TAR conformer simulations were analyzed using the same pipeline (*see associated scripts on Github*); those for which SciPy did not fit an unbound-state Gaussian were omitted. Contact frequency is presented as a fraction from 0 to 1 based on the fraction of bound-state frames that contain residue pairs within 15 Å of each other. Smaller cutoff distances implicated the same residues in binding as those shown in **Fig. 4B-C**. Dissociation constants calculated across the 15 well-sampled pairs are between ∼0.7 and 3.0 μM (based on normalization to SERF-rU30 affinity from simulations and experiments), which is remarkably consistent with the enhancement of binding affinity between SERF-(rU)30 and SERF-TAR measured in vitro (**Fig. S4C**).

### All-atom simulations in CAMPARI

#### Simulation setup and parameters

All-atom simulations of SERF were performed using the ABSINTH implicit solvent model and the CAMPARI Monte Carlo (MC) simulation engine (https://campari.sourceforge.net/) using the parameter set *abs3.5_opls.prm*. ABSINTH has been used to generate experimentally validated ensembles of a range of IDRs^104^. The *S. cerevisiae* SERF sequence (MARGNQRDLARQKNLKKQKDMAKNQKKSGDPKKRMESDAE ILRQKQAAADARREAEKLEKLKAEKTRR) with N- and C-terminal caps (acetyl and amide groups, respectively) was simulated in a spherical droplet with radius = 118 Å. The SERF simulations presented consist of frames merged from at least six independent simulations; three simulations used a ‘random’/extended starting conformation and three used a ‘helical’ starting conformation. The monovalent ion concentration (NaCl) was set at 0.05 M and was modeled explicitly. Each independent simulation reflects 50 million MC steps, of which the first 2.5 million were discarded as equilibration. The write-out frequency for accepted conformations was every 20,000 frames, such that the trajectories from each simulation contained >2,600 frames. Replicate trajectories from both starting conformations were merged to yield an ensemble of 14,250 total frames. See **Table S6** for the distribution of accepted moves. Unless otherwise specified, all simulations were performed at 300 K; it is important to note that the simulation temperature does not necessarily reflect a true 300 K, but rather was chosen to allow sufficient sampling of conformational space. ABSINTH/OPLS-AA parameters were used over ABSINTH/CHARMM36 in order to prevent aberrant chain collapse. For simulations of SERF with a fixed C-terminal helix, the AlphaFold2 (AF2)-predicted SERF structure was used to determine the boundary between N- and CTR of SERF. From the AF2 model, the C-terminal helix extends from residues 40 to 68; a modified PDB file containing only the helix between residues 40 and 68 was used as the structured input for simulations of SERF with a fixed C-terminal helix. In these simulations, the NTR of SERF and all side chains were allowed to sample.

#### Simulation analyses

CAMPARI simulations were analyzed using MDTraj and SOURSOP using custom Python scripts (see Github)^105,106^. Ensemble distributions for the analytical Flory random coil (AFRC) model were generated using the AFRC Google Colab notebook (https://colab.research.google.com/drive/1WHw8ous7IgcKd2LKYuJLeBTlkdEYoRAk?usp=sharing).

Paramagnetic relaxation enhancement (PRE) profiles were calculated from all-atom simulations as described in the references, but with modifications to account for the use of ^13^C direct-detect CON experiments for these measurements^43,107^. Profiles were calculated for both spin label locations (residues 10 and 63); because the simulated ensembles do not contain the covalent spin label, PRE calculations were performed using distance measurements from Cβ of the corresponding residue in the native SERF sequence. For reproducibility, the protocol is described in detail here. The ratio of cross-peak intensities between the paramagnetic (IP) and diamagnetic (ID) conditions is given by Equation n:

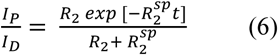

Where R2 is the intrinsic transverse relaxation rate (set to 8.1 s^-1^ for carbon nuclei), and *t* is the total duration of the inept delays in the CON experiment (*t* = 50 ms). *R2^sp^* is the PRE arising from the spin label that is experienced by the carbonyl ^13^C nucleus and is given by:

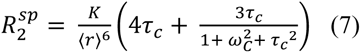

Where 〈*r*〉 is the ensemble-averaged distance from the Cβ of the ‘labeled’ residue to the carbonyl carbon of every residue in our simulations. *τ_c_* is the rotational correlation time estimated from the SERF molecular weight (7950 Da) based on reference to be ∼1 ns^108^. *ω_c_* is the Larmor frequency of the excitation nucleus, which is ^13^C (2*π* × 150.9 MHz) in CON experiments or ^1^H (2*π* × 600 MHz) in (HACA)CON experiments conducted at field strength of 14.1 T. *K* is an atom-specific parameter given by:

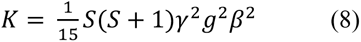

Where *S* is the spin angular momentum (S = ½ for ^1^H and ^13^C), *g* is the electronic g-factor (-2.002319, unitless), *β* is the Bohr magneton (9.274009994 × 10^-24^ J/T), and *γ* is the gyromagnetic ratio of the start nucleus (10.705 MHz/T for ^13^C or 26.751 MHz/T for ^1^H). Hence, KC = 7.78 × 10^-34^ cm^6^/s^2^ and KH = 1.23 × 10^-23^ cm^6^/s^2^. ^13^C direct-detect experiments for PRE used a modified ^13^C, ^15^N-CON pulse sequence that begins magnetization on Hα for improved sensitivity, so constants for ^1^H were used in calculating PRE profiles from simulations. To generate the PRE profile null model assuming a Gaussian distribution of mean squared distances, 〈*r*^)^〉, the following from Meng et al. was used ^43^:

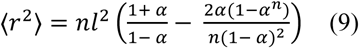

Where *n* is the number of residues between the site of the spin label and residue *i*, *l* is the monomer length (set to 3.8 Å), and *α* is the cosine of the bond-angle supplements given a freely rotating chain (set to 0.8). SAXS profiles were calculated from all-atom simulations using FoXS and then analyzed and visualized using BioXTAS RAW in the same manner as the experimental scattering data^109,110^. Structure visualization was performed using VMD or UCSF ChimeraX^111,112^.

## ACKNOWLEDGEMENTS

We thank Ke Wan in the Bardwell laboratory for protein purification. We are grateful to Dr. Debashish Sahu at the BioNMR Core of the University of Michigan and Dr. Bikash Sahoo for their help with NMR experiments. J. C. A. B. is an investigator at the Howard Hughes Medical Institute, which funded this work. The work in this study was supported by the following funding sources: NSF grant MCB-1932730 (to S. A. S.) for the NMR experiments, SIG S10 of the National Institutes of Health (NIH) under award number S10-OD028589 (to N. H. Y.) for SAXS experiments, and S10-OD030490 (to N. H. Y.) for the Wyatt SEC-MALS-DLS system. This project received funding from the (ERC) under the European Union’s Horizon 2020 research and innovation program (grant agreement No. 801728 PARAMIR to L.S.). Support for the development of native IM-MS technology for this project was provided by Agilent Technologies (to B. T. R.) and support from the National Institutes of Health to SAS (R01 DK121509) We also wish to thank Ms. Julia Fecko and Dr. Kevin E. W. Namitz for assistance at the X-ray Crystallography core at the Penn State Huck Institutes of the Life Sciences. R.M thanks Dr. Ben A. Meinen and Dr. Ursula H. Jakob for insightful discussions. E.T.U thanks Drs. Kacey Mersch, Tim Lohman, and Kathleen Hall for helpful discussions.

