## supplemental information for "Molecular insights into the interaction between a disordered protein and a folded RNA"

**This document includes:**

Figures S1 to S8
Tables S1 to S6

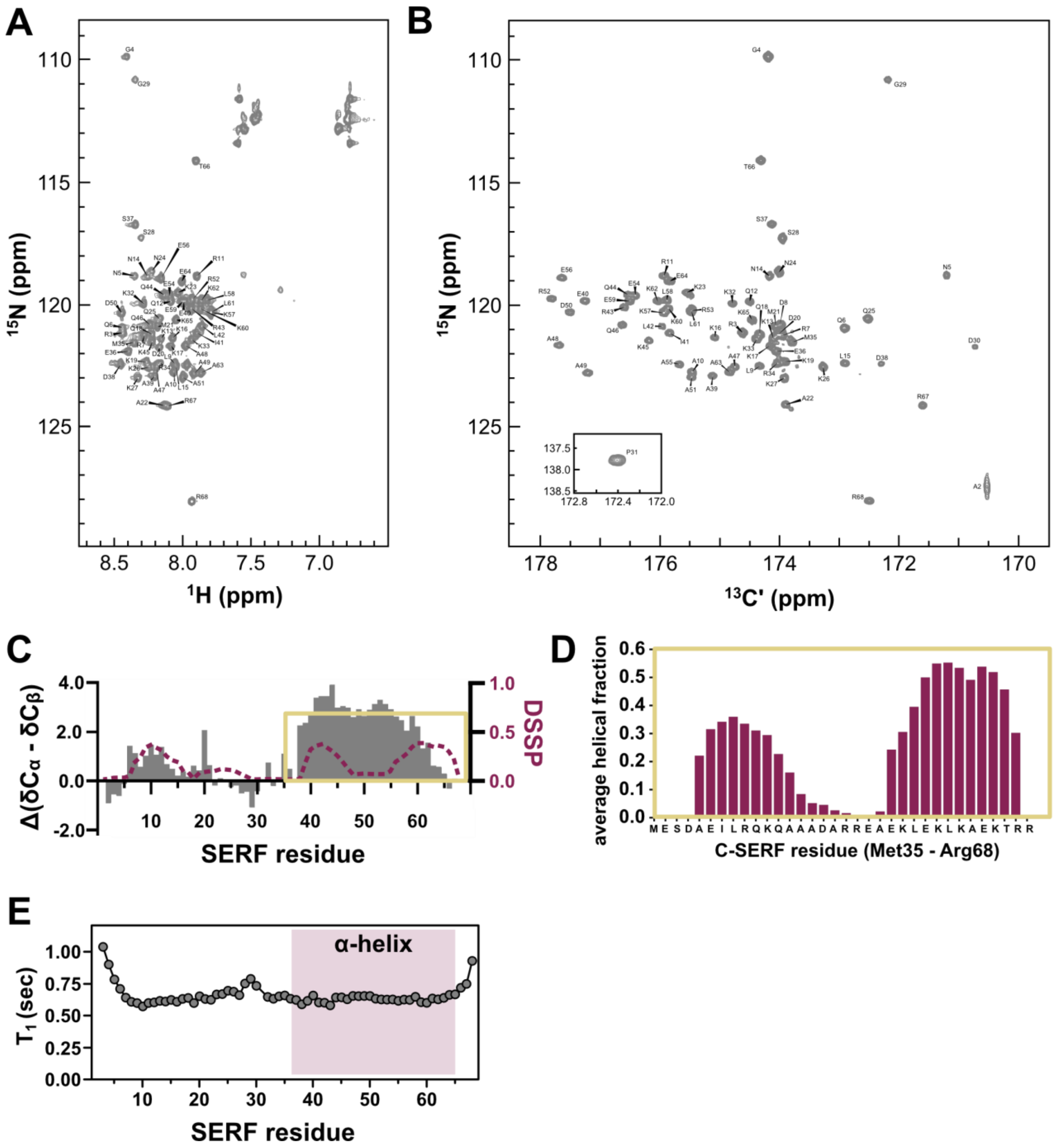


**Fig. S1: Local structural features of the SERF ensemble from NMR spectroscopy and all-atom simulations. (A)** $^{1}$H, $^{15}$N-HSQC spectrum with SERF assignments (see also Table S1). **(B)** $^{13}$C, $^{15}$N-CON spectrum of SERF with assignments (see also Table S1). **(C)** Plot of SERF secondary structure from side chain chemical shifts (positive values indicate α-helix; negative values indicate β-strand) (left axis). The DSSP scores for α-helical character averaged over all simulation frames are shown as a dashed line (right axis). The yellow rectangle shows the C-SERF helical region of interest. **(D)** Zoomed-in representation of DSSP scores for C-SERF helix from simulations. The breakpoint in the helix is centered around a stretch of charged residues (DARREAEK). **(E)** Plot of $T_1$ relaxation times per residue of SERF from $^{15}$N spin relaxation.

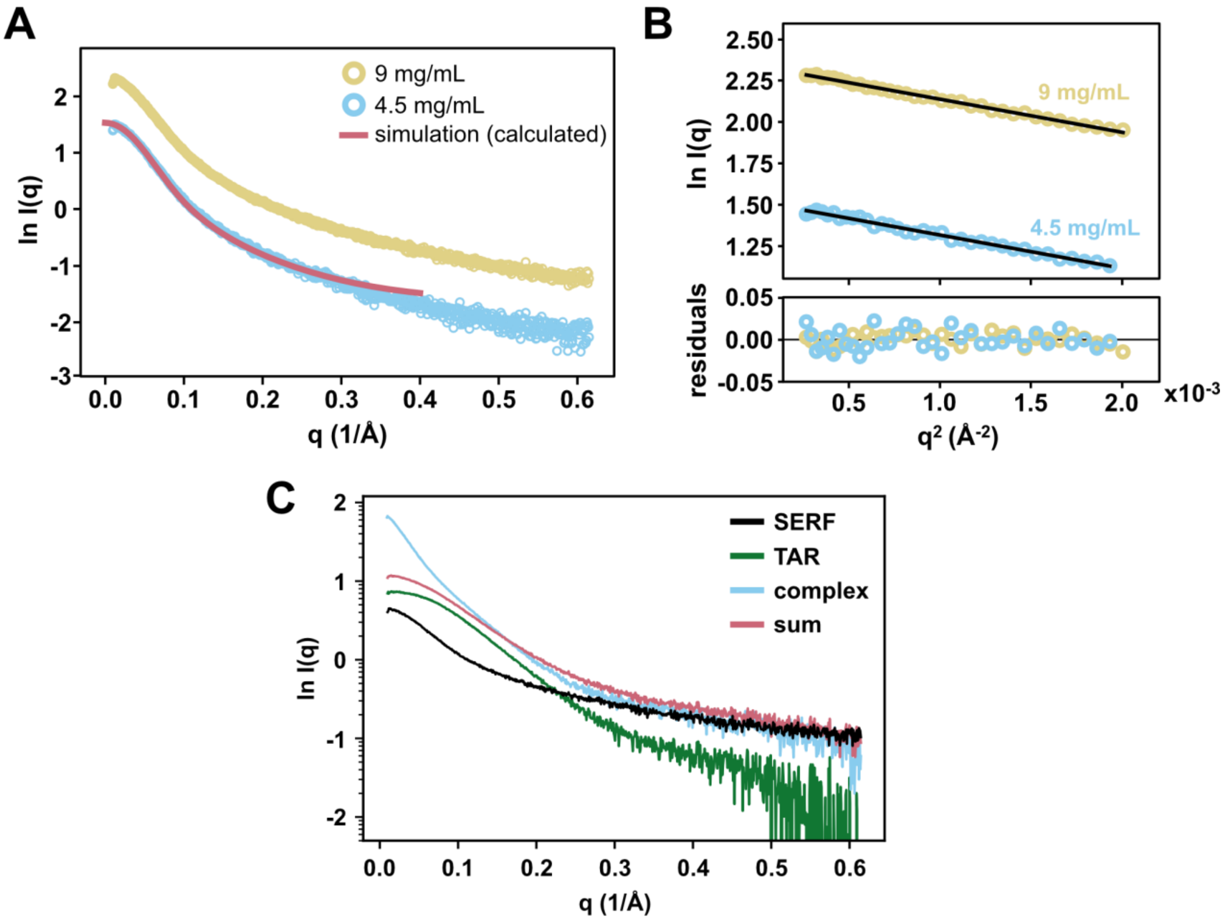


**Fig. S2: Global SERF ensemble characterization by SAXS and SV-AUC. (A)** Raw scattering curves for two concentrations of SERF shown with scaled SAXS profile calculated from all-atom simulations using FoXS(5). **(B)** Guinier transformation of SAXS data at two concentrations with linear fit to approximate radius of gyration. The small spread of residuals suggests adequate linear fitting in both cases. **(C)** Raw scattering curves for each SERF (black) and TAR (green) alone, the sum of their scattering profiles (pink), and the scattering curve measured for the complex (blue).

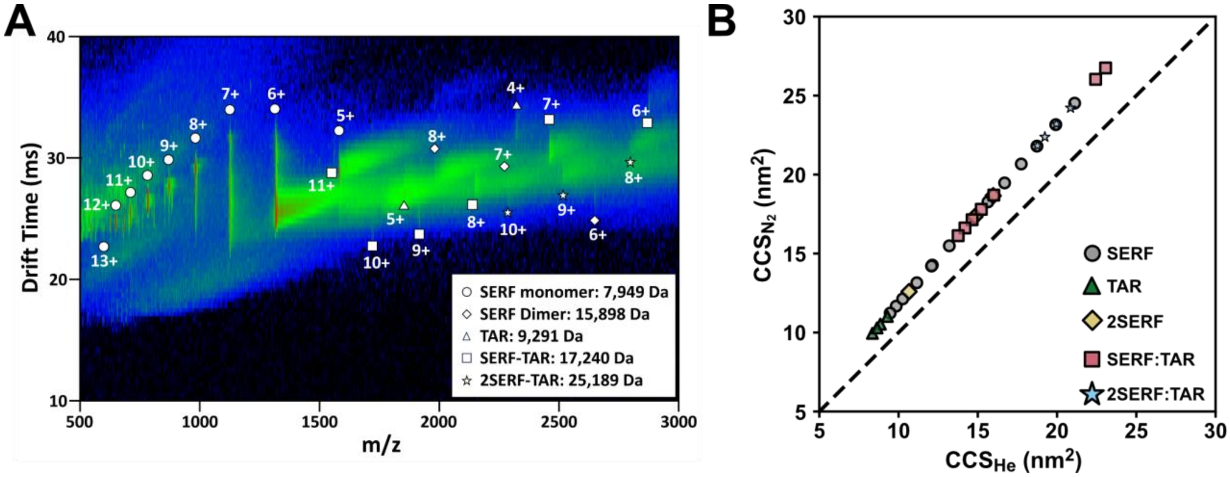


**Fig. S3: Ion-mobility mass spectrometry of SERF-TAR complexes. (A)** Plot of drift time as a function mass/charge ratio used to determine collision cross-section distributions. Each feature is assigned with the biomolecule(s) and ionization state it represents using the shapes given in the legend. The molecular weights for different species are given in the legend. **(B)** Plot of collision cross-sections from different carrier gases. The dashed line is the function y = x to guide the eye.

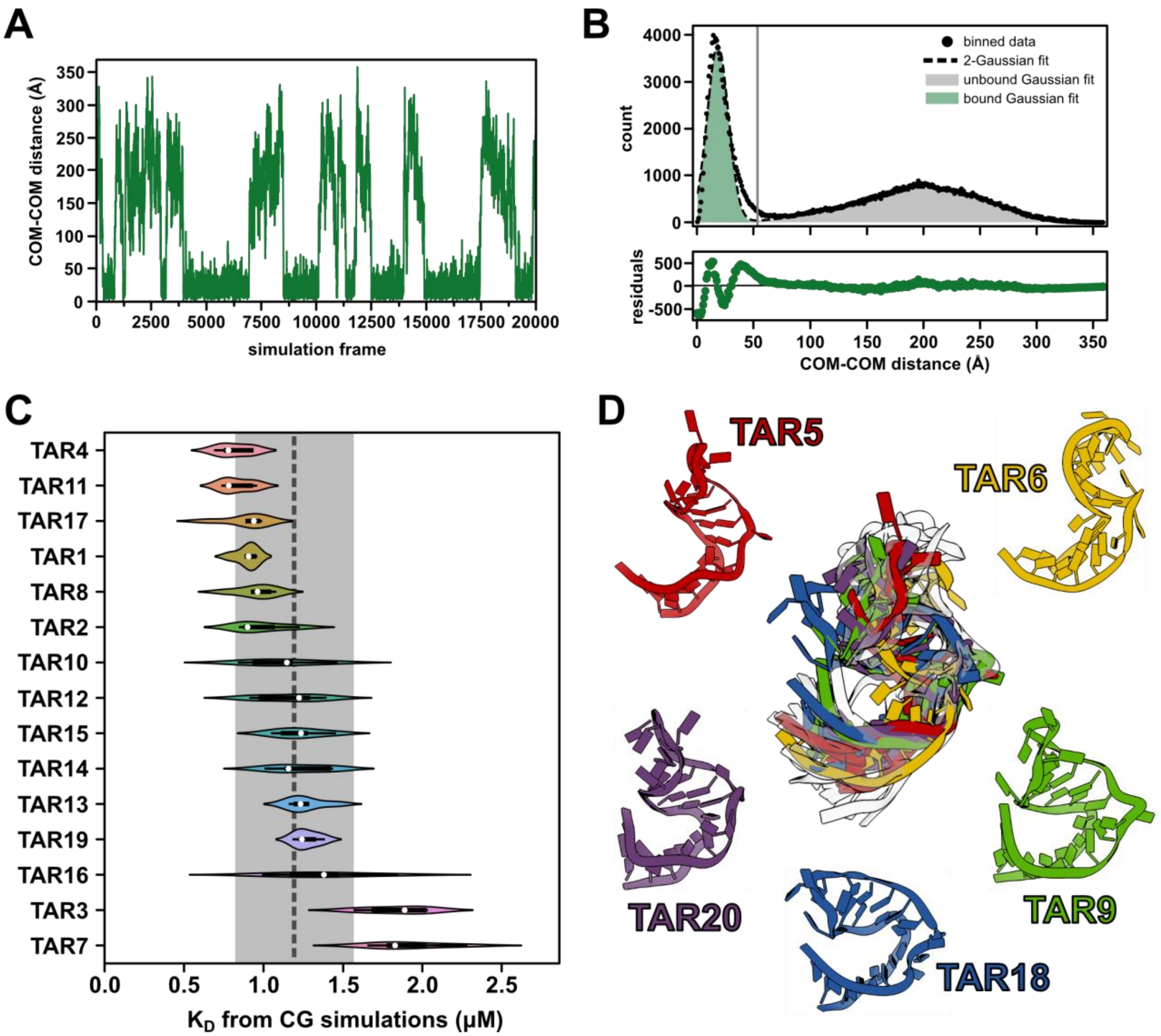


**Fig. S4: Monitoring 'SERF'-'TAR' binding by coarse-grained simulations using Mpipi.** **(A)** Trace of distances between each 'SERF' and 'TAR' over the course of the simulation measured from the center of mass (COM) of each molecule. **(B)** COM-COM distances from **(A)** presented as a distribution and fit with a 2-Gaussian model. The green (low COM-COM distance) and grey (high COM-COM distance) shading reflects 'bound' and 'unbound' frame assignments, respectively. The grey vertical line at 52.5 Å is the 'cutoff' distance (see *Methods*) that minimizes the overlap of the two sub-distributions. **(C)** Distribution of $K_D$ values across five replicates for each included TAR conformation ranked by average affinity. The vertical grey dashed line and shaded region represent the average and standard deviation of dissociation constants over all listed conformers. **(D)** Cartoon depiction of all 20 TAR conformations from PDB ID 1ANR aligned for visualization. Five colored TAR conformations that were omitted from average $K_D$ calculations are shown in the overlay and separately. See also Table S#.

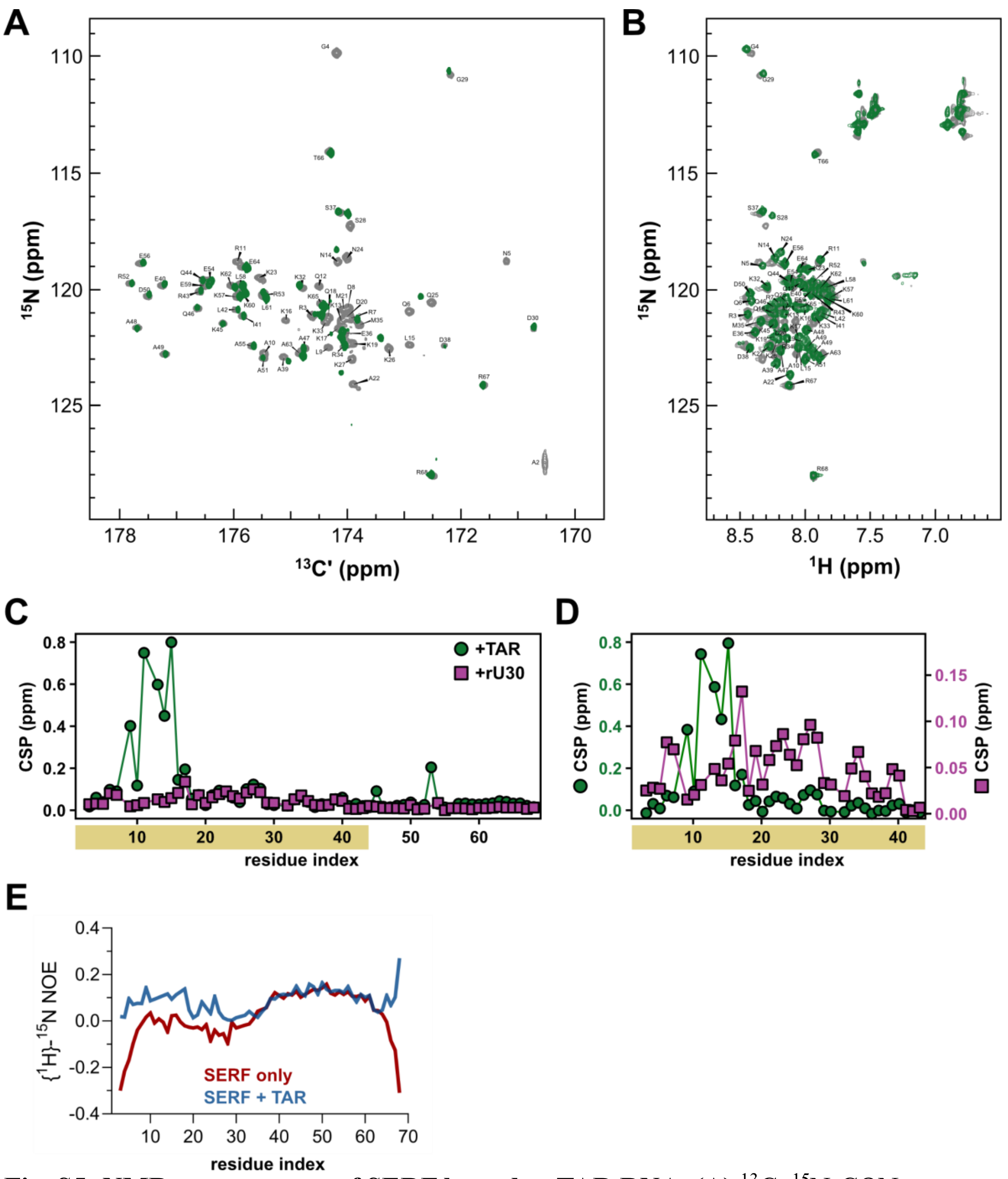


**Fig. S5: NMR spectroscopy of SERF bound to TAR RNA. (A)** $^{13}$C, $^{15}$N-CON spectra of unbound SERF (grey, from Fig. S2) and TAR-bound SERF (green). Significant line-broadening of resonances for residues that directly contact RNA precluded use of the CON for analysis of chemical shift perturbations upon RNA binding. Resonance assignments are shown for unbound SERF. **(B)** $^{1}$H, $^{15}$N-HSQC spectra of SERF alone (grey) and bound to TAR (green). The peak assignments shown are those for the bound state, which were used to generate the plots in **(C)** and **(D)**. **(C)** Plot of chemical shift perturbations (CSP) on SERF for binding to TAR RNA (green circles) or rU30 (magenta squares) on the same axis set. The yellow highlighting along the x-axis denotes the region of SERF that is examined more closely in **(D)**. **(D)** Re-scaled representation of CSP plot from **(C)** to reflect the differences in CSP magnitude for the different RNA binding partners. **(E)** Plot of heteronuclear NOEs per residue for the unbound (red) and RNA-bound SERF (blue).

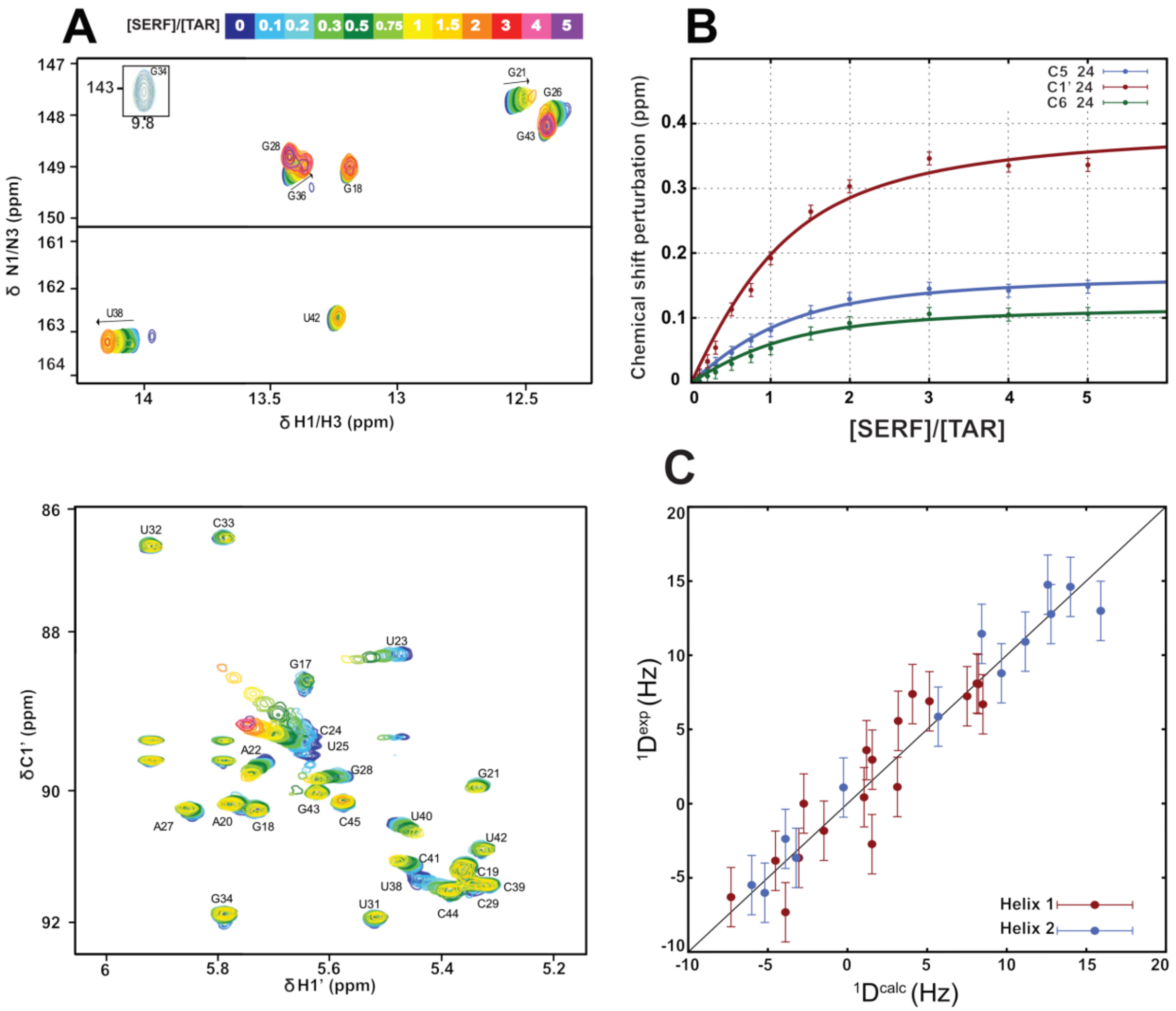

**Fig. S6: NMR spectroscopy of SERF-TAR interaction. (A)** (top) Imino and (bottom) C1' TROSY-HSQC spectra of isotopically enriched TAR showing chemical shift changes upon titrating with unlabeled SERF (molar ratio color coded according to the insert scale). **(D)** Chemical shift perturbations (in ppm) of the residue C24 of TAR with increasing [SERF]/[TAR] from C5H5 (red), C6H6 (green), and C1'H1' (blue). The lines correspond to the fit for a 1:1 complex with an apparent $K_d$ = 260 μM. Notably, this apparent $K_d$ differs from the $K_d$ obtained from the binding isotherm based on fluorescence anisotropy. The discrepancy in the two $K_d$ values might be due to the difference in protein and RNA concentrations present in the the two experiment types and possibly due to SERF-SERF interactions at high concentrations. Although the apparent $K_d$ value determined by NMR titrations is not directly used in this study, it highlights that the TAR:SERF complex is far from being saturated at a 1:1 ratio. **(C)** RDC reproduction of TAR assuming two independent ideal A-form helices. The associated alignment tensors are given in **Table S1**.

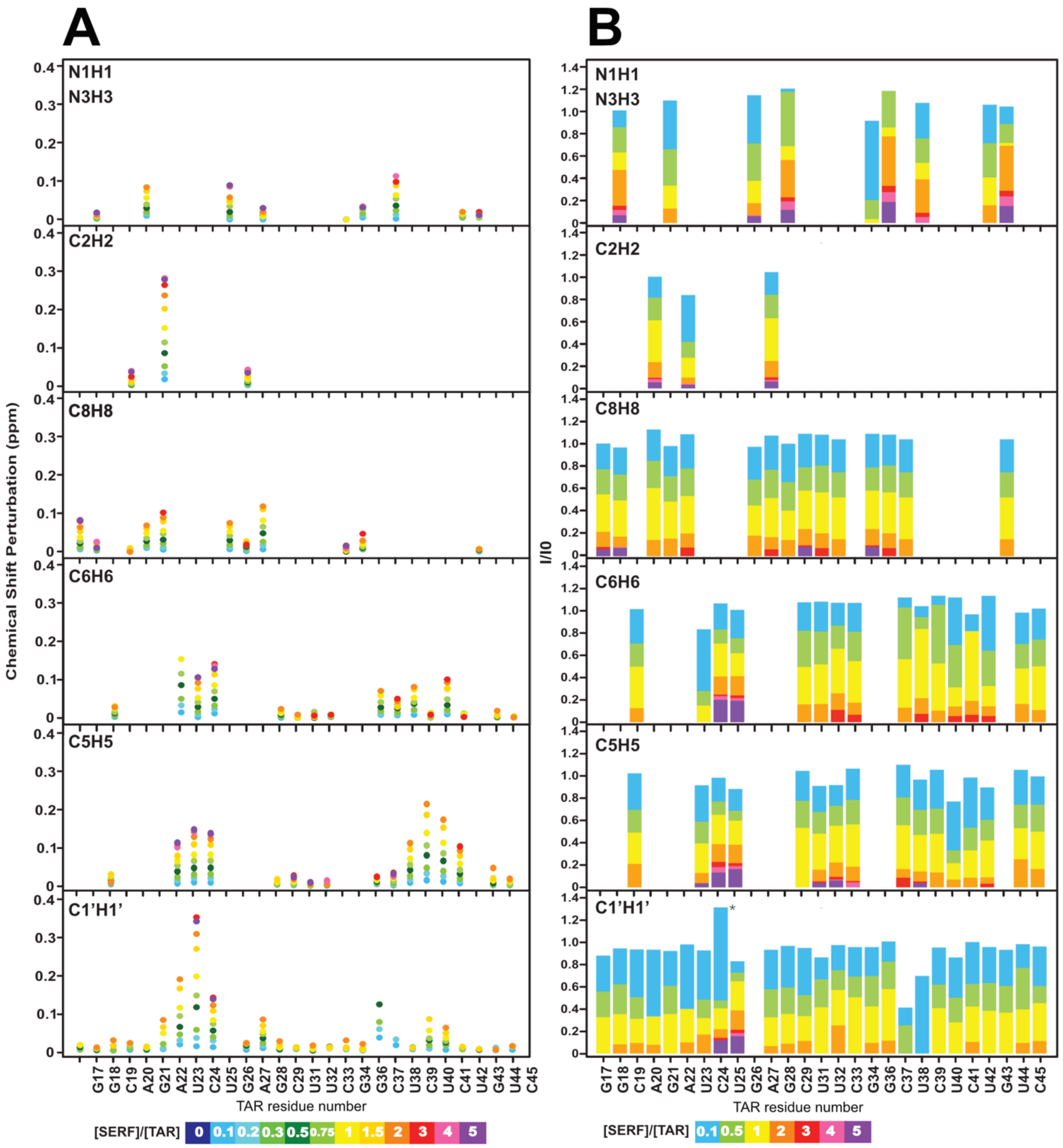


**Fig. S7: NMR titrations to probe the SERF-TAR interface.** (A) Chemical shift perturbations and (B) intensity ratios of TAR for N1H1, N3H3, C2H2, C6H6, C8H8, C5H5, and C1'H1' spin pairs along the titration (**Fig 5 and S6**). Molar ratio color coded according to the insert scale. The star (*) indicates a higher uncertainty for signal highly overlapping in the TAR alone spectrum.

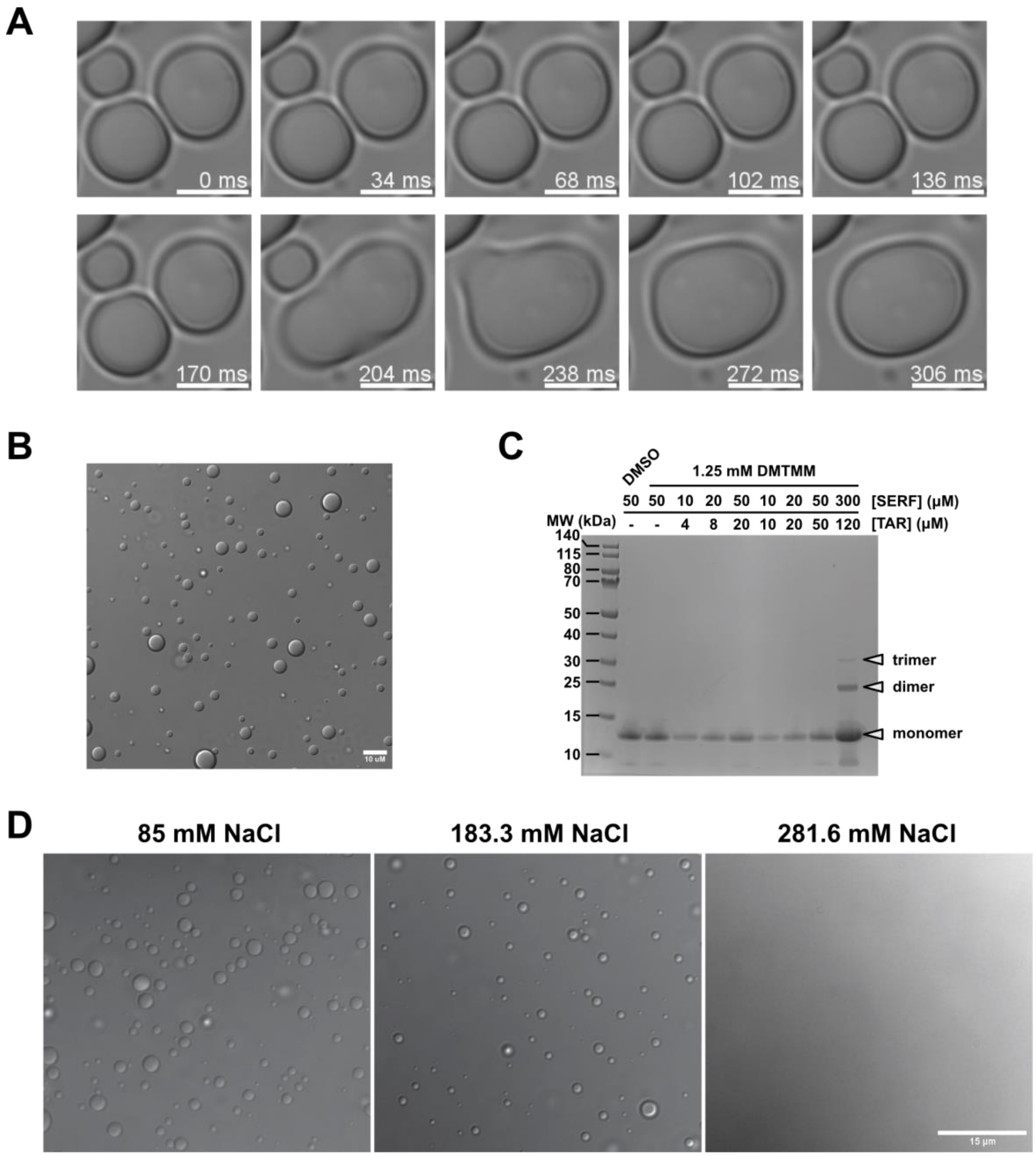


**Fig. S8: SERF and TAR undergo concentration-dependent assembly and phase separation.** **(A)** Time-resolved DIC images of SERF-TAR droplets (in 10% PEG buffer) demonstrate their liquid-like behavior via droplet fusion. **(B)** DIC image of SERF-TAR droplets composed of 125 μM TAR and 312.5 μM SERF in 0% PEG. **(C)** SDS-PAGE showing DMTMM-crosslinked SERF species in the presence or absence of TAR RNA. Higher-order SERF species are observed only at high concentrations of SERF and TAR. **(D)** DIC images of SERF-TAR droplets (in 10% PEG buffer) showing sensitivity to NaCl concentrations.

| Residue # | Amino acid | Hα (ppm) | Cα (ppm) | Cβ (ppm) | CO (ppm) | N (ppm) |
|---|---|---|---|---|---|---|
| 1 | M | 3.94 | 54.36 | 32.64 | 170.56 | -- |
| 2 | A | 4.31 | 50.93 | 18.47 | 174.63 | 127.49 |
| 3 | R | 4.27 | 55.39 | 30.25 | 174.20 | 121.13 |
| 4 | G | 3.89 | 44.60 | n/a | 171.22 | 109.85 |
| 5 | N | 4.61 | 52.40 | 38.17 | 172.93 | 118.80 |
| 6 | Q | 4.16 | 56.17 | 28.20 | 173.95 | 120.95 |
| 7 | R | 4.17 | 56.24 | 29.69 | 173.98 | 121.28 |
| 8 | D | 2.61 | 53.91 | 40.15 | 174.34 | 120.80 |
| 9 | L | 4.12 | 55.55 | 41.22 | 175.46 | 122.51 |
| 10 | A | 4.16 | 52.73 | 17.83 | 175.94 | 122.75 |
| 11 | R | 4.16 | 56.29 | 29.62 | 174.50 | 118.81 |
| 12 | Q | 4.15 | 55.93 | 28.32 | 174.10 | 119.86 |
| 13 | K | 4.15 | 56.49 | 32.17 | 174.17 | 121.39 |
| 14 | N | 4.58 | 52.96 | 37.85 | 172.91 | 118.84 |
| 15 | L | 4.19 | 55.06 | 41.46 | 175.09 | 122.39 |
| 16 | K | 4.15 | 56.03 | 32.09 | 174.15 | 121.34 |
| 17 | K | 4.18 | 56.02 | 32.15 | 174.33 | 121.69 |
| 18 | Q | 4.21 | 55.66 | 28.58 | 173.91 | 121.21 |
| 19 | K | 4.17 | 56.17 | 32.23 | 174.03 | 122.32 |
| 20 | D | *n.d.* | 56.01 | 40.24 | 174.07 | 121.21 |
| 21 | M | 4.33 | 55.13 | 31.32 | 173.90 | 120.88 |
| 22 | A | 4.20 | 51.50 | 18.18 | 175.55 | 124.09 |
| 23 | K | 4.16 | 56.00 | 32.15 | 174.02 | 119.50 |
| 24 | N | 4.61 | 52.63 | 38.01 | 172.54 | 118.62 |
| 25 | Q | 4.23 | 55.18 | 28.76 | 173.28 | 120.56 |
| 26 | K | 4.22 | 55.66 | 32.33 | 173.92 | 122.54 |
| 27 | K | 4.29 | 55.43 | 32.38 | 173.96 | 122.99 |
| 28 | S | 4.36 | 57.65 | 63.20 | 172.19 | 117.27 |
| 29 | G | 3.88 | 44.23 | - | 170.74 | 110.79 |
| 30 | D | 4.79 | 51.41 | 40.58 | 172.42 | 121.72 |
| 31 | P | 4.32 | 62.93 | 31.47 | 174.79 | 137.76 |
| 32 | K | 4.15 | 56.07 | 31.82 | 174.39 | 119.95 |
| 33 | K | 4.22 | 55.40 | 32.12 | 174.03 | 121.40 |
| 34 | R | 4.21 | 55.74 | 29.91 | 173.79 | 122.40 |
| 35 | M | 4.33 | 55.38 | 31.43 | 174.05 | 121.54 |
| 36 | E | 4.25 | 56.31 | 29.45 | 174.13 | 121.90 |
| 37 | S | 4.37 | 57.86 | 63.21 | 172.30 | 116.70 |
| 38 | D | 4.43 | 55.41 | 39.78 | 175.12 | 122.41 |

| Residue # | Amino acid | Hα (ppm) | Cα (ppm) | Cβ (ppm) | CO (ppm) | N (ppm) |
|---|---|---|---|---|---|---|
| 39 | A | 4.05 | 53.99 | 17.78 | 177.25 | 122.92 |
| 40 | E | 4.13 | 57.75 | 28.43 | 175.84 | 119.83 |
| 41 | I | 3.73 | 63.23 | 37.00 | 175.96 | 121.15 |
| 42 | L | 4.06 | 56.99 | 40.77 | 176.60 | 120.88 |
| 43 | R | 4.07 | 58.35 | 29.37 | 176.52 | 120.09 |
| 44 | Q | 4.06 | 57.72 | 27.57 | 176.18 | 119.59 |
| 45 | K | 4.07 | 58.38 | 31.80 | 176.62 | 121.47 |
| 46 | Q | 4.05 | 57.14 | 27.67 | 174.75 | 120.81 |
| 47 | A | 4.19 | 53.71 | 17.23 | 177.67 | 122.56 |
| 48 | A | 4.18 | 53.55 | 17.35 | 177.20 | 121.66 |
| 49 | A | 4.09 | 53.90 | 17.36 | 177.49 | 122.79 |
| 50 | D | 4.11 | 55.55 | 39.56 | 175.46 | 120.30 |
| 51 | A | 4.14 | 53.82 | 17.25 | 177.80 | 122.97 |
| 52 | R | 4.11 | 57.64 | 29.11 | 175.46 | 119.74 |
| 53 | R | 4.09 | 58.06 | 29.43 | 176.40 | 120.17 |
| 54 | E | 4.06 | 57.87 | 28.44 | 175.67 | 119.62 |
| 55 | A | 4.13 | 53.99 | 17.41 | 177.63 | 122.44 |
| 56 | E | 4.02 | 57.94 | 28.76 | 175.93 | 118.90 |
| 57 | K | 4.00 | 57.88 | 31.74 | 175.88 | 120.33 |
| 58 | L | 4.07 | 56.26 | 41.00 | 176.50 | 119.82 |
| 59 | E | 4.01 | 57.83 | 28.69 | 175.86 | 119.84 |
| 60 | K | 4.10 | 57.62 | 31.75 | 175.48 | 120.14 |
| 61 | L | 4.14 | 55.83 | 41.34 | 176.04 | 120.28 |
| 62 | K | 4.06 | 57.05 | 31.98 | 174.84 | 119.85 |
| 63 | A | 4.17 | 52.40 | 18.12 | 175.83 | 122.78 |
| 64 | E | 4.14 | 56.42 | 29.29 | 174.46 | 119.04 |
| 65 | K | 4.25 | 55.99 | 32.12 | 174.32 | 120.62 |
| 66 | T | 4.16 | 61.43 | 69.61 | 171.62 | 114.10 |
| 67 | R | 4.29 | 55.41 | 30.01 | 172.51 | 124.12 |

**Table S1: Chemical shift assignments for SERF from $^{13}C$ direct-detect NMR experiments.** Atoms with missing assignments denoted by -- are those for which we do not expect to observe a resonance (i.e., we cannot detect the amide nitrogen for N-terminal residue in $^{13}C$-detected experiments; and Gly residues do not have Cβ atoms). Aside from these, only one atom is missing a chemical shift assignment (Hα from Asp20). n.d. - not determined.

**Table S2: SERF ensemble dimensions as described by radius of gyration.**

| system | method | $R_G$ (Å) | error (type) |
|---|---|---|---|
| *sequence-based predictions* | | | |
| SERF | AFRC (random coil) | 20.8 | N/A |
| | ALBATROSS | 24.8 | N/A |
| *X-ray scattering (experiments)* | | | |
| SERF | Guinier approx. | 24.9 | 0.1 (SE fit) |
| | EOM | 25.2 | N/A |
| | MFF | 23.9 | 0.12 (SE fit) |
| *simulations* | | | |
| SERF | CAMPARI (all-atom) | 25.9 | 0.8 (SD of replicate means) |
| | mPiPi (coarse-grained) | 25.4 | 0.1 (SD of conformer means) |
| SERF (+ $r(U)_{29}$)* | mPiPi | 22.9 | |
| SERF (+ TAR)* | mPiPi | 22.1 | 0.2 (SD of conformer means) |

*the $R_G$ calculations for SERF in complex were performed on the protein chain only; the RNA chain was ignored. N/A – not applicable.

**Acronyms:** AFRC – analytical Flory random coil(1); ALBATROSS – deep learning-based predictor(2); EOM – ensemble optimization method(3); MFF – molecular form factor(4); SE – standard error; SD – standard deviation.

**Table S3: Dissociation constants from experiments and simulations.**

| Protein | RNA | $K_D$ (μM)† | error‡‡ |
|---|---|---|---|
| *experiments* | | | |
| SERF | TAR | 0.67 | 0.04 |
| SERF | $r(U)_{30}$ | 1.9 | 0.2 |
| *coarse-grained simulations* | | | |
| SERF | TAR | 1.2 | 0.1 |
| SERF | $r(U)_{29}$ | 4.1 | 0.6 |
| $SERF_{1-34}$ | TAR | 1.1 | 0.1 |
| $SERF_{1-34}$ | $r(U)_{29}$ | 3.6 | 0.7 |

†Dissociation constants from coarse-grained simulations are not absolute values; a correction factor of 10 was introduced into the analysis workflow for easier comparison between experiments and simulations.

‡From experiments, the reported error is the standard deviation of the fit from the covariance matrix; the error from simulations is the standard error of the mean calculated from $K_D$ values across independent replicates.

**Table S4: Masses and collision cross-sections (CCSs) of all SERF, TAR, and SERF-TAR complex ions detected by IM-MS with nitrogen or helium carrier gas.**

| ID | Ion Mass | Ion Charge (z) | CCS, $N_2$ ($nm^2$) | CCS, $N_2$ S.D. | CCS, He ($nm^2$) | CCS, He S.D. |
|---|---|---|---|---|---|---|
| **SERF** | 7949.4 (+/- 0.2 Da) | 13 | 24.52 | 0.02 | 21.11 | 0.02 |
| | | 12 | 23.16 | 0.05 | 19.92 | 0.04 |
| | | 11 | 21.80 | 0.01 | 18.73 | 0.01 |
| | | 10 | 20.67 | 0.00 | 17.74 | 0.00 |
| | | 9 | 19.47 | 0.01 | 16.69 | 0.01 |
| | | 8 | 18.27 | 0.01 | 15.64 | 0.01 |
| | | 7 | 13.15 | 0.02 | 11.15 | 0.02 |
| | | 7 | 14.27 | 0.04 | 12.13 | 0.03 |
| | | 7 | 15.49 | 0.01 | 13.20 | 0.01 |
| | | 7 | 17.32 | 0.01 | 14.80 | 0.01 |
| | | 6 | 11.67 | 0.03 | 9.85 | 0.02 |
| | | 6 | 12.13 | 0.08 | 10.26 | 0.07 |
| | | 6 | 14.21 | 0.10 | 12.08 | 0.09 |
| | | 5 | 11.23 | 0.01 | 9.47 | 0.01 |
| **2SERF** | 15898 (+/- 1 Da) | 8 | 17.34 | 0.03 | 14.82 | 0.03 |
| | | 8 | 18.66 | 0.09 | 15.97 | 0.08 |
| | | 6 | 12.59 | 0.10 | 10.66 | 0.08 |
| **TAR** | 9290.7 (+/- 0.4 Da) | 5 | 10.53 | 0.01 | 8.85 | 0.01 |
| | | 5 | 11.02 | 0.06 | 9.29 | 0.05 |
| | | 4 | 9.95 | 0.02 | 8.35 | 0.02 |
| | | 4 | 10.29 | 0.06 | 8.64 | 0.05 |
| **SERF-TAR** | 17240 (+/- 2 Da) | 11 | 26.03 | 0.19 | 22.43 | 0.16 |
| | | 11 | 26.75 | 0.18 | 23.06 | 0.16 |
| | | 10 | 18.70 | 0.07 | 16.01 | 0.06 |
| | | 9 | 17.80 | 0.02 | 15.22 | 0.02 |
| | | 8 | 17.14 | 0.06 | 14.64 | 0.05 |
| | | 7 | 16.62 | 0.05 | 14.19 | 0.04 |
| | | 6 | 16.13 | 0.04 | 13.76 | 0.03 |
| **2SERF-TAR** | 25189 (+/- 1 Da) | 11 | 23.20 | 0.12 | 19.96 | 0.10 |
| | | 11 | 24.22 | 0.10 | 20.85 | 0.09 |
| | | 10 | 22.38 | 0.11 | 19.23 | 0.09 |
| | | 9 | 21.83 | 0.09 | 18.76 | 0.08 |

**Table S5: Alignment tensor derived from the analysis of residual dipolar couplings (RDCs) measured on TAR alone and in a non-saturated TAR-SERF 1:1 complex.**

| System | Helix | Aa ($10^{-4}$) | Ar ($10^{-4}$) | $\alpha$ (°) | $\beta$ (°) | $\gamma$ (°) | Scalar Product |
|---|---|---|---|---|---|---|---|
| TAR | 1 | 4.79 | 0.27 | 35 | 38 | -76 | 0.24 |
| TAR | 2 | 8.19 | 0.53 | -50 | 17 | 44 | 0.24 |
| TAR-SERF | 1 | 3.25 | 1.55 | -134 | 170 | -67 | 0.82 |
| TAR-SERF | 2 | 8.34 | 0.59 | -27 | 168 | 19 | 0.82 |

Alignment tensors are expressed using their axial (Aa) and rhombic (Ar) components, as well as three Euler angles defining their orientations.

**Table S6: Acquisition parameters for protein NMR experiments**

| Experiments | Time domain data size (points) | # scans | sweep widths (ppm) | | | Transmitter frequency offsets (ppm) | | |
|---|---|---|---|---|---|---|---|---|
| | | | $^1H$ | $^{15}N$ | $^{13}C$ | $^1H$ | $^{15}N$ | $^{13}C$ |
| $^1H^N$-start CON-based $^{15}N$- $T_1$ | 1024 ($^{13}C$) $\times$ 256 ($^{15}N$) $\times$ 13 | 16 | n/a | 35 | 12 | n/a | 122.5 | 174 |
| $^1H^N$-start CON-based $^{15}N$- $T_2$ | 1024 ($^{13}C$) $\times$ 256 ($^{15}N$) $\times$ 16 | 16 | n/a | 35 | 12 | n/a | 122.5 | 174 |
| PRE- CON | 1024 ($^{13}C$) $\times$ 256 ($^{15}N$) | 128 | n/a | 35 | 12 | n/a | 122.5 | 174 |
| PRE- (HACA)CON | 1024 ($^{13}C$) $\times$ 256 ($^{15}N$) | 128 | n/a | 35 | 12 | n/a | 122.5 | 174 |
| (HACA)N(CA)CON | 1024 ($^{13}C$) $\times$ 64 ($^{15}N$) $\times$ 128 ($^{15}N$) | 8 | n/a | 35 | 12 | n/a | 122.5 | 174 |
| (HACA)N(CA)NCO | 1024 ($^{13}C$) $\times$ 64 ($^{15}N$) $\times$ 128 ($^{15}N$) | 16 | n/a | 35 | 12 | n/a | 122.5 | 174 |
| (CC)(CO)N | 1024 ($^{13}C$) $\times$ 64 ($^{15}N$) $\times$ 128 ($^{13}C$) | 8 | n/a | 35 | 12, 80 | n/a | 122.5 | 174 |
| H(CC)(CO)N | 1024 ($^{13}C$) $\times$ 64 ($^1H$) $\times$ 128 ($^{15}N$) | 8 | 14 | 35 | 12 | 4.69 | 122.5 | 174 |
| HNCO | 1024 ($^1H$) $\times$ 64 ($^{15}N$) $\times$ 128 ($^{13}C$) | 16 | 16 | 35 | 11 | 4.672 | 122.5 | 174 |
| HN(CA)CO | 1024 ($^1H$) $\times$ 92 ($^{15}N$) $\times$ 60 ($^{13}C$) | 16 | 16 | 35 | 11 | 4.672 | 117 | 176.8 |
| HNCACB | 1024 ($^1H$) $\times$ 64 ($^{15}N$) $\times$ 128 ($^{13}C$) | 8 | 16 | 35 | 80 | 4.673 | 117 | 43 |
| CBCA(CO)NH | 1024 ($^1H$) $\times$ 64 ($^{15}N$) $\times$ 128 ($^{13}C$) | 8 | 16 | 35 | 80 | 4.673 | 117 | 43 |
| H(CC)(CO)NH | 2048 ($^1H$) $\times$ 64 ($^{15}N$) $\times$ 128 ($^1H$) | 16 | 16 | 35 | n/a | 4.673 | 117 | n/a |
| $^1H$-$^{15}N$ HSQC-based $^{15}N$- $T_1$ | 1024 ($^1H$) $\times$ 256 ($^{15}N$) $\times$ 13 | 16 | 16 | 35 | n/a | 4.673 | 117 | n/a |
| $^1H$-$^{15}N$ HSQC-based $^{15}N$- $T_2$ | 1024 ($^1H$) $\times$ 256 ($^{15}N$) $\times$ 16 | 16 | 16 | 35 | n/a | 4.7 | 117 | n/a |

n/a - not applicable
